# Essential genes for *Haemophilus parainfluenzae* survival and biofilm growth

**DOI:** 10.1101/2024.03.31.587483

**Authors:** Thais H. de Palma, Chris Powers, Morgan J. McPartland, Jessica Mark Welch, Matthew Ramsey

## Abstract

*Haemophilus parainfluenzae* (*Hp*) is a Gram-negative, pleomorphic rod, highly prevalent and abundant as a commensal in the human oral cavity, and an infrequent extraoral opportunistic pathogen. *Hp* occupies multiple niches in the oral cavity, including the tongue dorsum, keratinized gingiva, and the supragingival plaque biofilm. As a member of the HACEK group, *Hp* is also known to cause infective endocarditis. Additionally, case reports have identified *Hp* as the causative agent of meningitis, septic arthritis, chronic osteomyelitis, septicemia, and a variety of other infectious diseases. Little is known about how *Hp* interacts with its neighbors in the healthy biofilm nor about its mechanisms of pathogenesis as an extraoral opportunistic pathogen. To address these unknowns, we identified the essential genomes of two *Hp* strains and the conditionally essential genes for their growth in *in vitro* biofilms aerobically and anaerobically. Using transposon insertion sequencing (TnSeq) with a highly saturated *mariner* transposon library in two strains, the ATCC33392 type-strain (*Hp* 392) and a commensal oral isolate EL1 (*Hp* EL1), we show that the essential genome of *Hp* 392 and *Hp* EL1 is composed of 395 and 384 genes, respectively. The core essential genome, consisting of 341 essential genes conserved between both strains, was composed of genes associated with genetic information processing, carbohydrate, protein, and energy metabolism. We also identified conditionally essential genes for aerobic and anaerobic biofilm growth, which were associated with carbohydrate and energy metabolism in both strains of *Hp*. Additionally, RNAseq analysis determined that most genes upregulated during anaerobic growth are not essential for *Hp* 392 anaerobic biofilm survival. The completion of this library and analysis under these conditions gives us a foundational insight into the basic biology of *H. parainfluenzae* in differing oxygen conditions, similar to its *in vivo* oral habitat. Further, the creation of this library presents a valuable tool for further investigation into conditionally essential genes for an organism that lives in close contact with many microbial species in the human oral habitat.

## Introduction

The oral cavity harbors a diverse microbial community with more than 700 species of bacteria capable of colonizing the mouth (Aas et al., 2005; Chen et al., 2010; Dewhirst et al., 2010). Oral microbes are exposed to mechanical and physical disturbances such as oral hygiene practices, mastication, salivary flow and others (Mark Welch et al., 2019). Thus, microbes must attach to host substrates and/or each other forming an attached community known as a biofilm to persist in the mouth (Kolenbrander et al., 2006; Marsh, 2005). The biofilm formed on the tooth surface above the gum line is known as supragingival plaque (SUPP). Among the factors that affect the composition and organization of many biofilms, oxygen stands out as a major factor (Ahn and Burne, 2007). While the mouth is largely aerobic, oxygen availability in SUPP varies. The periphery of the biofilm is rich in oxygen, while its interior is mostly anaerobic and rich in CO_2_ (Wessel et al., 2014). This oxygen gradient within SUPP allows for aerobes, obligate anaerobes, and facultative anaerobes to coexist (Diaz et al., 2006). One facultative anaerobe commonly found in SUPP is *Haemophilus parainfluenzae* (*Hp*), a Gram-negative bacterium. *Hp* is also found in high abundance in other niches of the oral cavity, such as the tongue dorsum, keratinized gingiva, and saliva (Kilian and Schiott, 1975; Mark Welch et al., 2019, 2016; Utter et al., 2020), demonstrating that *Hp* is a generalist in the mouth (Mark Welch et al., 2019).

*Hp* is commonly isolated from the mouth of healthy individuals (Caselli et al., 2020) and is not typically associated with oral disease. However, elsewhere in the body, *Hp* can act as an opportunistic pathogen. *Hp* is a member of the HACEK group (*Haemophilus, Aggregatibacter, Cardiobacterium hominis, Eikenella corrodens*, and *Kingella*), which is made up of bacterial species and genera that commonly colonize the oropharynx and can cause infective endocarditis (Sen Yew et al., 2014). Besides endocarditis, *Hp* has also been infrequently associated with other diseases including meningitis (Bachman, 1975; Black et al., 1988), septicemia (Oill et al., 1979), pleural effusion (Cremades et al., 2011), urethritis (Sturm, 1986), prosthetic joint infection (Bailey et al., 2011), and respiratory infections such as pneumonia (Oill et al., 1979; Pillai et al., 2000), pharyngitis and epiglottitis (Oill et al., 1979), and chronic obstructive pulmonary disease (Smith et al., 1976).

To fully understand what is required for *Hp* to exist as a key member of the healthy oral microbiome and an infrequent extraoral opportunistic pathogen, it is useful to know the minimal set of genes that *Hp* needs to survive. A current method to determine essential genes is to use a high-throughput method known as transposon sequencing (TnSeq) (Van Opijnen et al., 2009). This technique has been used to assess bacteria fitness under different conditions (Klein et al., 2015; Narayanan et al., 2017; Turner et al., 2014) and has also been applied to identify the essential genome of other bacterial species that colonize the oral cavity and the upper respiratory tract such as *Streptococcus mutans* (Walker and Shields, 2022), *Aggregatibacter actinomycetemcomitans* (Narayanan et al., 2017), *Porphyromonas gingivalis* (Klein et al., 2015), *Treponema denticola* (Yang et al., 2008), and *Haemophilus influenzae* (Gawronski et al., 2009). In addition to TnSeq, transcriptomes can shed light on the adaptation of an organism to its environment and allow for hypotheses generation for mechanisms that may be essential for fitness in that environment. For example, transcriptome measurement via RNAseq has been used to identify the influence of oxygen in gene expression in other microorganisms (Vergara-Irigaray et al., 2014) and has proven to be a very sensitive and comprehensive method for characterizing bacterial transcriptomes (Croucher and Thomson, 2010).

Here, we describe the essential genes necessary for *Hp* survival and the conditionally essential genes required for *Hp* to survive in a biofilm aerobically and anaerobically. Given the pronounced genomic variation among *Hp*, (Kerr et al., 1993; Watts et al., 2021) we characterized essential genes in two different *Hp* strain backgrounds. Comparison of essential genes in both strains allowed us to identify the genes that were conserved and essential both in *Hp* 392 and *Hp* EL1 -named here the “absolute essential genome of *Hp*” and differentiate them from strain-specific essential genes. Our results reveal that 20% and 21% of the genes present in *Hp* 392 and *Hp* EL1, respectively, are essential for survival *in vitro* and that a large majority of essential genes identified are conserved between strains. We then further describe the conditionally essential genes necessary for both strains to survive in *in vitro* biofilms aerobically and anaerobically, along with the genes that are differentially expressed in those conditions measured by RNAseq. Identifying the essential genome of *Hp* and the conditionally essential genes for biofilm growth aerobically and anaerobically provide a basic framework of likely genes necessary for *in vivo* colonization and present a valuable tool for the study of *Hp* in both oral niches as a commensal and extraoral niches as an opportunistic pathogen.

## Results

### Transposon library development and sequencing

We created saturating *mariner* transposon libraries via methods we have used previously (Narayanan et al., 2017), conjugating a delivery vector into two *Hp* strains, the ATCC3392 type-strain and the oral isolate EL1 (de Palma et al., 2023), generating ∼200,000 and ∼100,000 individual mutants respectively. Similar to previous methods (Narayanan et al., 2017; Turner et al., 2015, 2014), we generated and sequenced Illumina libraries to map *mariner* insertion sites in each library. Libraries were generated in triplicate for each experiment. We identified 195,115 unique insertions present in at least one of three replicates in *Hp* 392 and 144,946 in *Hp* EL1, resulting in a coverage of approximately one insertion at every 11 and 15 bp, respectively, indicating that the mutant pools were well saturating and randomly distributed (Fig. 1A, 1B).

**Figure 1.**
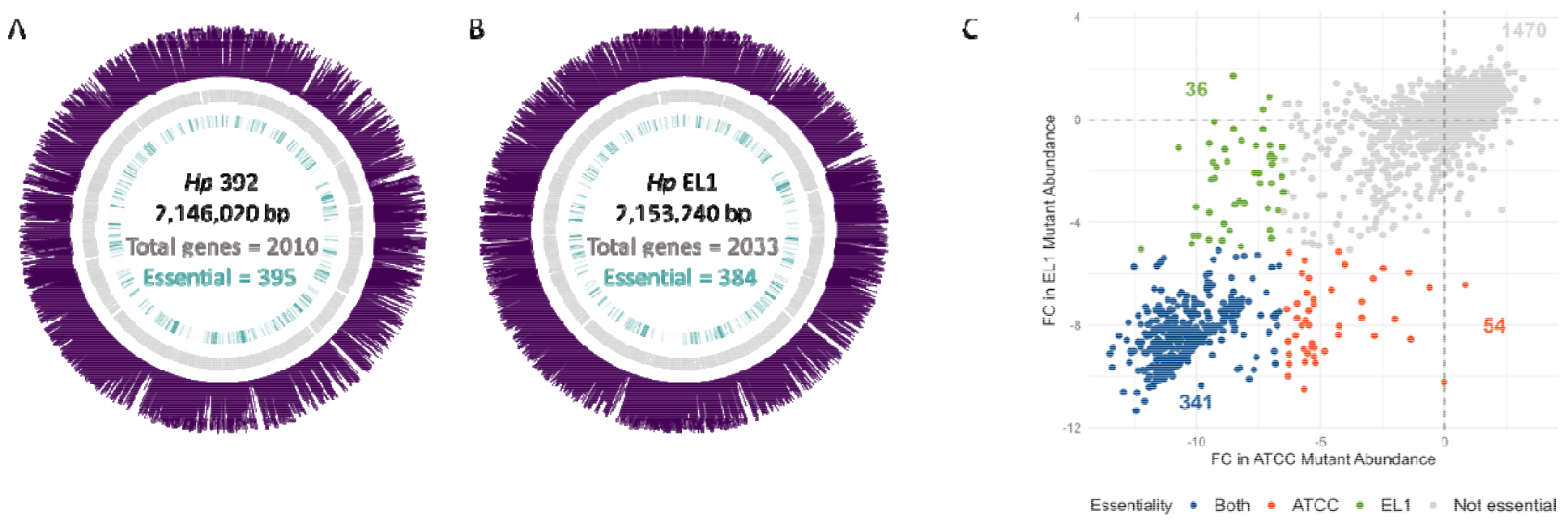
The essential genome of *Hp* 392 (A) and *Hp* EL1(B). The outer (purple) ring indicates transposon insertions, next inward (gray) represents open reading frames, and the inner ring (green) indicates essential genes. **(C)** shows the log2 fold change (FC) in mutant abundance for *Hp* 392 (x axis) versus EL1 (y axis). Each point corresponds to an ortholog and genes < 200 bp were excluded from the analysis.

### The essential genome for *H. parainfluenzae*

Analysis of insertion adjacent sequences determined which genes were not represented in the mutant library, *i*.*e*., genes that render *Hp* unfit for growth and thus are absolutely essential for *in vitro* aerobic conditions on nutrient-rich medium used for library selection. We then used a Monte Carlo-based approach as previously described (Lewin et al., 2019; Narayanan et al., 2017; Turner et al., 2015) to verify essential genes. Genes considered essential are expected to have zero, or significantly fewer, transposon insertions compared to non-essential genes, reflecting their inability to grow when these genes are interrupted and their absence from the outgrown library. We observed that 395 (21%) and 384 (20%) genes are essential in *Hp* 392 and *Hp* EL1, respectively, and that most essential genes had zero transposon insertions (Fig. S1). We then grouped these genes into their respective KEGG categories (Kanehisa, 2019; Kanehisa et al., 2023; Kanehisa and Goto, 2000) and determined that most of the essential genes were, as expected, involved in key functions including genetic information processing and common energy metabolism pathways (Fig. 2). 102 and 109 genes that were < 200 bp in length were excluded from the analysis for *Hp* 392 and *Hp* EL1, respectively, due to low insertion coverage, leading to unacceptably high variability for the determination of gene essentiality.

**Figure 2.**
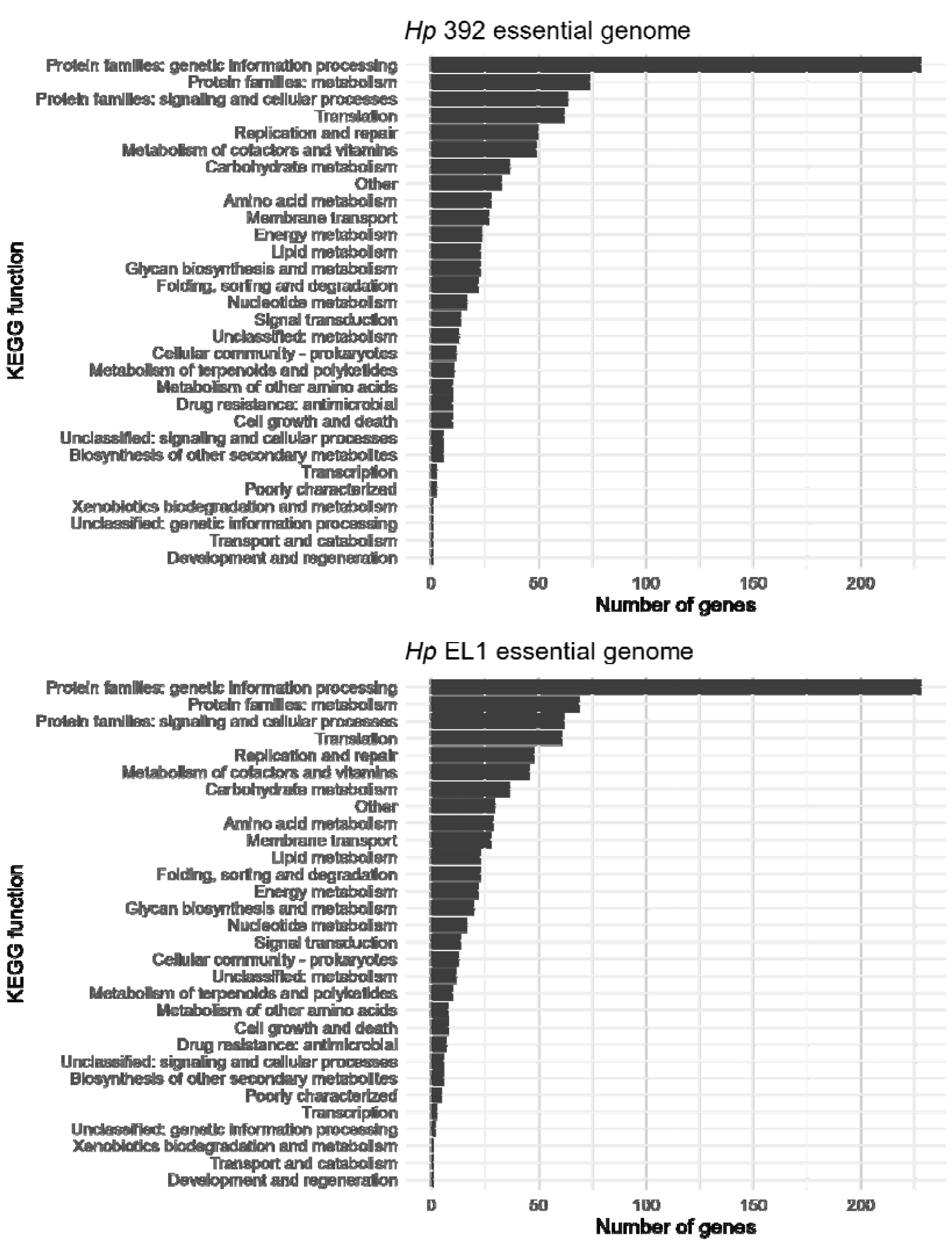
KEGG group assignments for absolute essential genes in *Hp*. The essential genome of *Hp* 392 (A) and *Hp* EL1(B) grouped into KEGG categories. Genes < 200 bp were not included in the analysis. Most essential genes are involved in central cell functional processes including genetic information processing and common energy metabolism pathways.

### Core essential genes for *H. parainfluenzae*

Comparison of the type-strain *Hp* 392 genome to the commensal oral isolate EL1 revealed that 1835 genes are shared between both strains, representing 91% and 90% of their respective genomes (Table S7). Of these 1835 shared genes, we found that 341 were absolutely essential for survival *in vitro* (Fig. 1C). We then compared these 341 core essential genes via Anvi’o (Eren et al., 2021) to 16 *Hp* closed genomes available in the NCBI database (Fig. 3) and found that all 341 genes were conserved between all 16 genomes. These data indicate that the core essential genes detected in our transposon screen under laboratory conditions are also conserved in natural populations. This is an interesting observation, given the genomic plasticity of *Hp* (Kerr et al., 1993; Watts et al., 2021).

**Figure 3.**
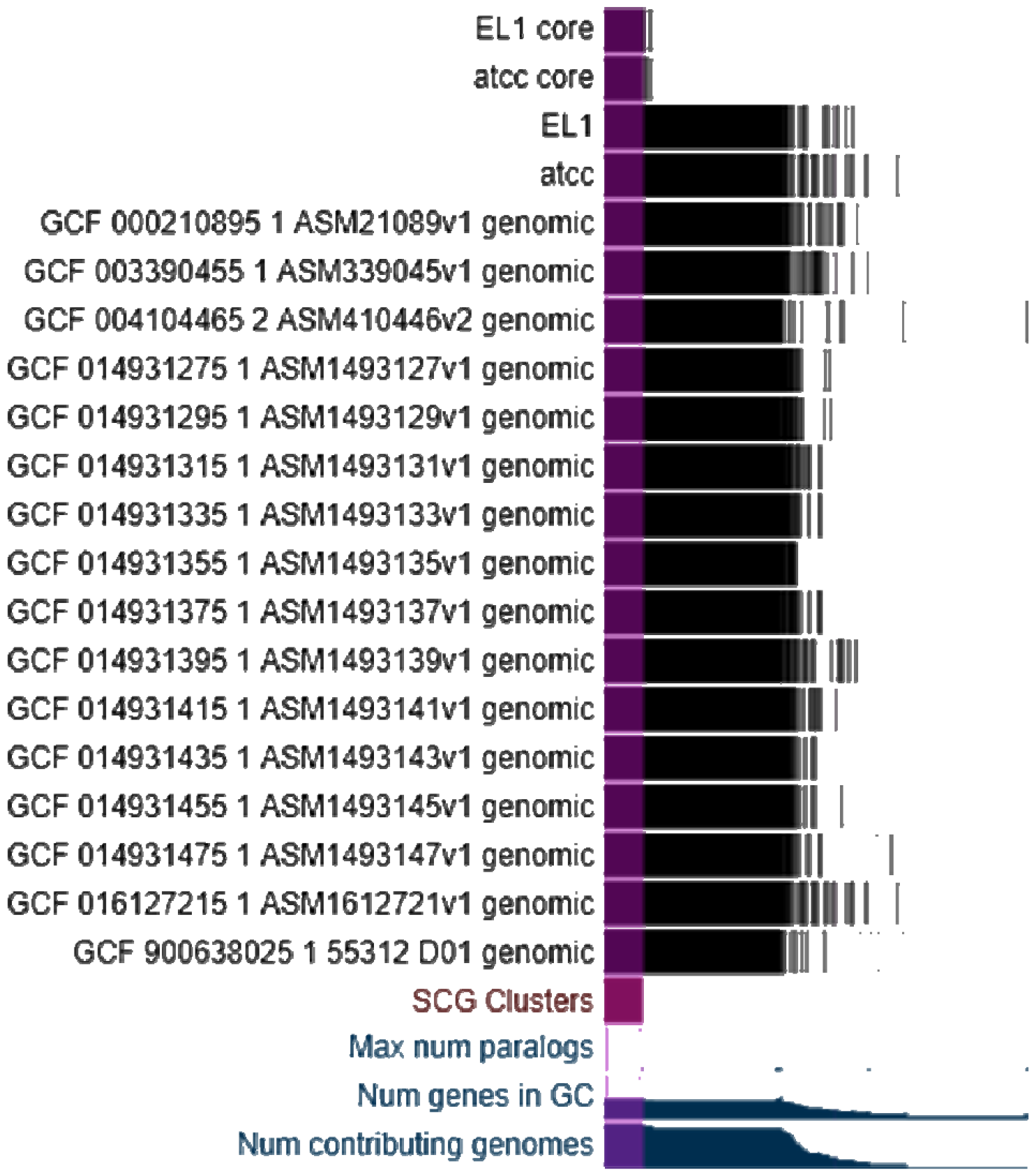
The core essential genome of *Hp*. Comparison of 18 genomes of *Hp* and the core essential genome of *Hp* 392 and *Hp* EL1 (top two rows) with 16 available *Hp* genomes. Purple indicates the core essential genome, *i*.*e*., the TnSeq identified absolute essential genes that are conserved in all species.

### *H. parainfluenzae* conditionally essential genes for *in vitro* aerobic and anaerobic biofilm growth

To identify genes conditionally essential *in vitro* for either aerobic or anaerobic biofilm growth, *Hp* were grown using a colony biofilm model (Anderl et al., 2000) on a permeable membrane on solid medium as we have done previously (Perera et al., 2021) and incubated either in the presence or absence of oxygen in 5% CO_2_ for 24h. Colony biofilms were scraped from the membranes, and DNA extraction and sequencing were performed to assess which genes were “conditionally” essential for growth in aerobic and anaerobic biofilms for both strains (Table 1). We next classified the conditionally essential genes based on their predicted KEGG functions (Fig. 4), observing that they are largely involved with carbohydrate and energy metabolism as anticipated. A full list of all genes and their predicted KEGG functions is presented in tables S3 and S4.

**Table 1.** Number of conditionally essential genes for *Hp* 392 and *Hp* EL1aerobically and anaerobically. Previously essential genes and those < 200 bp are not shown.

| Strain | Aerobic biofilm | Anaerobic biofilm | Aerobic and anaerobic biofilm |
| --- | --- | --- | --- |
| *Hp 392* | 73 | 55 | 50 |
| *Hp EL1* | 119 | 121 | 108 |
| **Both** | 36 | 28 | 27 |

**Figure 4.**
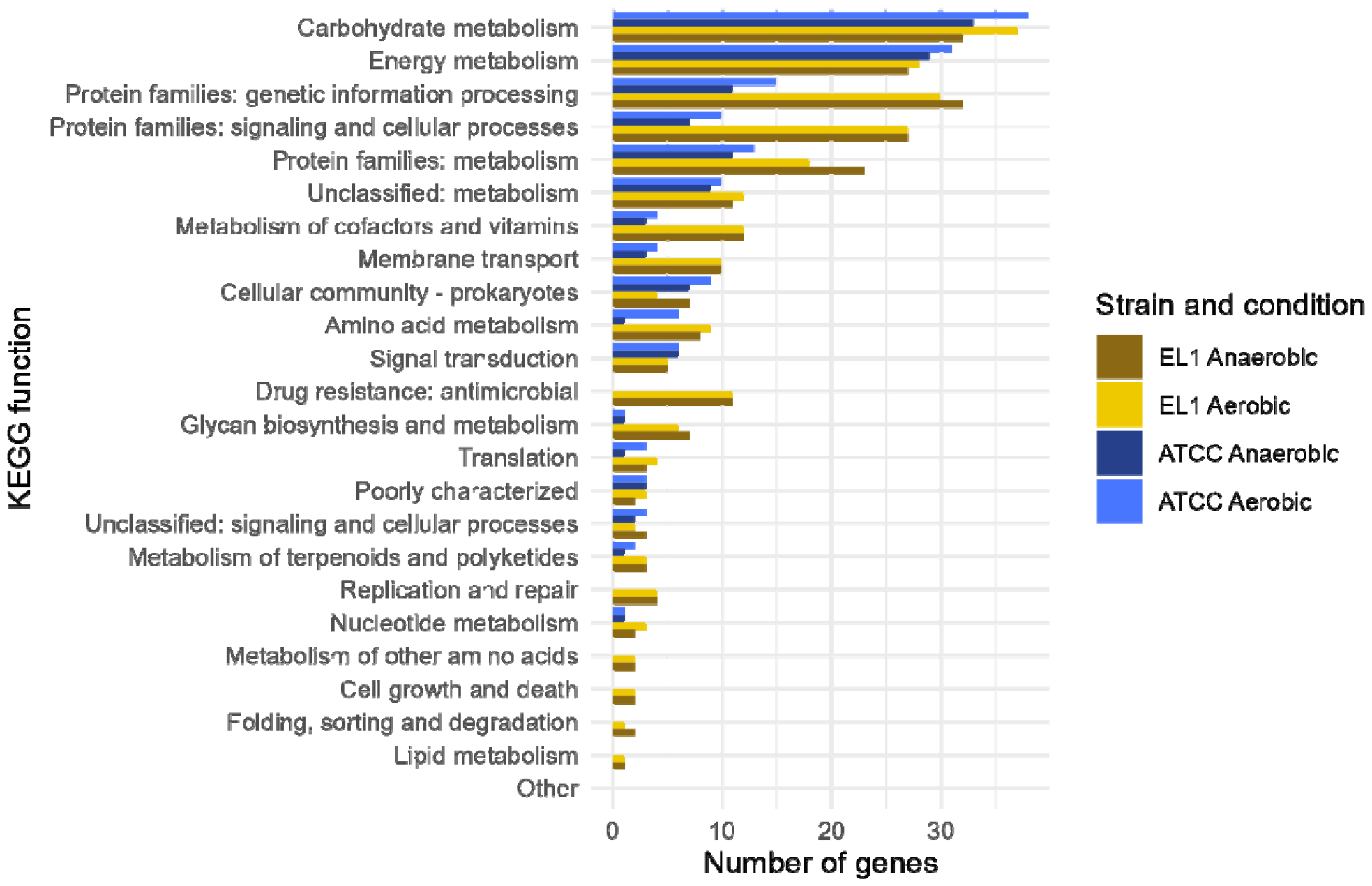
Genes essential for aerobic and anaerobic growth in a colony biofilm model for *Hp* 392 and *Hp* EL1. Conditionally essential genes for aerobic and anaerobic growth in *in vitro* biofilms in *Hp* 392 (blue), and *Hp* EL1 (yellow) grouped into higher KEGG functions.

### The *Hp* 392 anaerobic transcriptome

To determine which genes are differentially regulated in aerobic vs. anaerobic biofilm conditions, we assayed *Hp* 392 grown aerobically and anaerobically in a colony biofilm assay via RNAseq. This analysis revealed 167 genes that were significantly differentially expressed (either up or downregulated ≥ 2-fold) between the two conditions (Table S5A and S5B). Ninety-eight differentially expressed genes (DEGs) were upregulated in anaerobic biofilm conditions and 69 downregulated (Table S5A and S5B, respectively). Genes upregulated were largely involved in genetic information processing (26%), metabolism (11%), and signaling and cellular processes (10%). The downregulated genes were involved in signaling and cellular processes (38%), genetic information processing (19%), and membrane transport (13%). Overall transcriptome responses were largely unremarkable, and we saw broad similarity between both environments, likely due to the transiently anaerobic nature of even aerobic biofilms.

### Comparison of *in vitro* anaerobic biofilm transcriptome to conditionally essential genes for anaerobic biofilm growth in *Hp* 392

To identify whether genes that are induced under anaerobic biofilm growth conditions are also conditionally essential for anaerobic biofilm growth, we compared the conditionally essential genes (TnSeq data) to differentially expressed genes (RNAseq data) (Fig. 5). We observed that out of the 98 genes upregulated in anaerobic conditions, only three (Murein DD-endopeptidase MepM, LPS-core synthesis glycosyltransferase PM0509, and Bacterial ribosome SSU maturation protein RimP) were also considered essential for anaerobic biofilm growth. Moreover, one downregulated gene (Phosphoenolpyruvate carboxykinase [ATP]) was also essential. Taken together, these results indicate that there is almost no overlap between gene essentiality and differential expression, reminiscent of previous findings in *Pseudomonas aeruginosa* (Turner et al., 2014).

**Figure 5.**
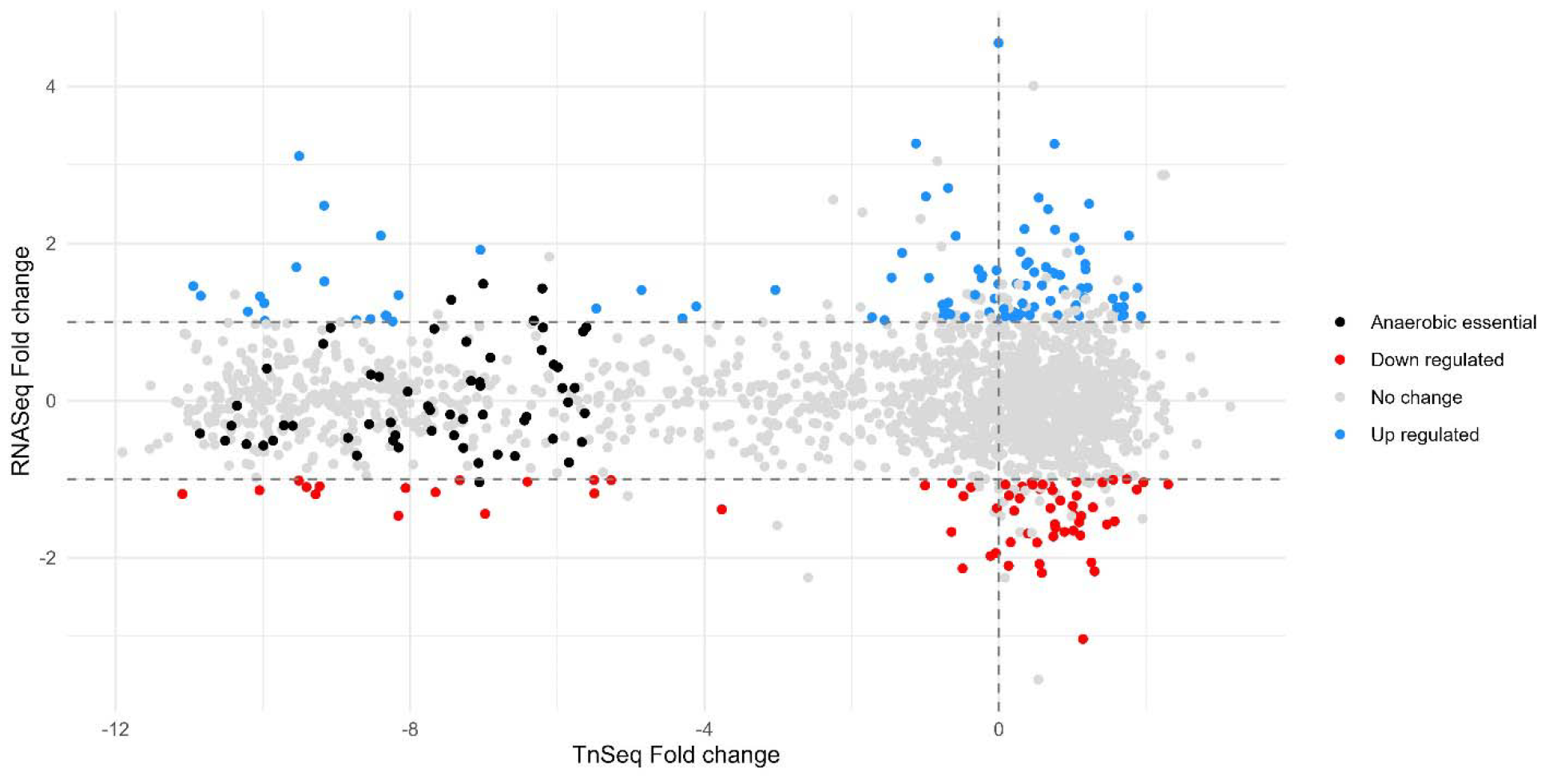
The majority of *Hp* 392 genes conditionally essential for anaerobic biofilm growth are not differentially regulated between aerobic and anaerobic conditions. 98 genes, indicated in blue, were significantly upregulated (p<= 0.05, fold change >=2) in anaerobic compared to aerobic conditions, while 67 genes, indicated in red, were significantly down regulated (p<= 0.05, fold change <=-2). Genes uniquely essential for colony growth under anaerobic conditions are indicated in black with only three total being significantly differentially expressed.

## Materials and Methods

### Strains and growth conditions

Strains and plasmids used are listed in Table S1. *Hp* ATCC 33392 and *Hp* EL1 (de Palma et al., 2023) were used to build *mariner* mutant libraries. *Hp* strains were grown in brain heart infusion (BHI) supplemented with 15 μg/ml of nicotinamide adenine dinucleotide (NAD), 15 μg/ml of hemin, and 5% yeast extract (BHIYE-HP). For agar plates, the medium was supplemented with 1.6% agar. Cultures were incubated at 37 °C, in 5% CO_2_ overnight unless otherwise specified. *Escherichia coli* strains were grown in Luria-Bertani (LB) broth and agar supplemented with 0.3 mM of diaminopimelate (DAP) (for the MFD*pir* conjugation host) and incubated at 37 °C overnight.

### *H. parainfluenzae* transposon library construction

*mariner* mutant pools were developed as previously described (Narayanan et al., 2017) with some modifications. Individual colonies of *Hp* 392 and *Hp* EL1 were inoculated into 5 mL of BHIYE-HP and incubated overnight at 37 °C, in 5% CO_2_. Single colonies of *E. coli* MFD*pir*, harboring the *mariner* transposon delivery plasmid pMR361-K, were inoculated into 5 mL of LB and incubated at 37 °C, shaken at 200RPM. Next, five independent conjugations for *Hp* 392 and four conjugations for EL1 were made. For that, samples were washed with PBS, and OD_600_ was adjusted to 1 for *Hp* strains and 0.1 *E. coli* MFD*pir. Hp* cultures were then incubated at 46 °C for 6 min. Next, 10 µl of *Hp* and 10 µl of *E. coli* MFD*pir* were combined and plated on BHIYE-HP supplemented with 0.3 mM of DAP and incubated at 37 °C, 5% CO_2_ overnight. Colonies were scraped from the plate and resuspended in 2 mL of BHIYE diluted in 50% glycerol. Cells were then plated on the counter-selection plate containing BHIYE-HP supplemented with 0.1 mM isopropyl-β-D-thiogalactopyranoside (IPTG), 40 mg/ml kanamycin, and 30 mg/ml nalidixic acid (to select against *E. coli* donor strains). Counterselection plates were incubated at 37 °C, 5% CO_2_ for 48h. Colonies were then scraped and resuspended in 2ml of BHIYE plus 50% glycerol. The total number of cells in each conjugation was calculated via CFU counting. Approximately 40,000 cells were grown in eight plates for each of the five conjugations of *Hp* 392, and 25,000 in five plates for each of the four conjugations of *Hp* EL1, totaling approximately 200,000 colonies for *Hp* 329, and more than 100,000 for EL1. Plates were incubated for 48h, and colonies were scraped, homogenized, and aliquoted into cryogenic storage medium with 25% glycerol. Mutant pool stocks were stored at -80 °C.

### Identification of essential genes

approximately 2.5×10^6^ cells of *Hp* 392 and *Hp* EL1 from the mutant pools’ stocks were inoculated into 3 ml of BHIYE-HP broth and incubated for 16 hours at 37 °C, 5% CO_2_. Cultures were centrifuged for 1 minute at 14,000 g, and cells were collected for DNA extraction using the Epicentre® kit following the manufacturer’s instructions. Samples were lysed using a Mini-Beadbeater (Biospec) in 2 mL vials preloaded with Lysing Matrix B (MP Biomedicals). Library growth was performed in biological triplicates using three different aliquot tubes to start respective cultures.

### The core genome of *H. parainfluenzae*

Anvi’o (Eren et al., 2021; Hyatt et al., 2010) was used to identify the core genome and the core essential genome of *Hp*. To do so, the genomes of *Hp* 392 and *Hp* EL1, and the 341 essential genes shared by both strains were aligned to 16 closed *Hp* genomes available at the NCBI database (Table S5). We used scripts available on GitHub (https://github.com/mramsey01/TnSeq-analysis-1). Briefly, modified FASTA files were loaded into the Anvi’o default pangenome pipeline for alignment and comparison.

### TnSeq Illumina Library preparation

to locate *mariner* insertion sites in the genome, Illumina libraries were prepared largely as described previously (Turner et al., 2015, 2014). DNA was extracted and sheared to 400 bp average length (peak power of 175.0 and 200 cycles/Burst) using a Covaris S220 Focused-ultrasonicator. Small and large fragments (< 100 bp or > 700 bp) of DNA were removed using KAPA beads according to the manufacturer’s instructions. Next, two rounds of PCR were performed using primers (Table S2) targeting the transposon and adding the Illumina adapters for sequencing similar to previous methods (Turner et al., 2015, 2014). Libraries were submitted to the Rhode Island Genomics and Sequencing Center (RIGSC) for quantification (qPCR) and quality control (Bioanalyzer). Libraries were then sent to SeqCenter and sequenced on an Illumina NextSeq 550 1-by-75 single-end run at the Microbial Genome Sequencing Center (MIGS) Pittsburgh, PA.

### *H. parainfluenzae* conditionally essential genes for *in vitro* aerobic and anaerobic biofilm growth

*Hp* 392 and *Hp* EL1 mutant pools were grown in 3 ml of BHIYE-HP broth and incubated for 16 hours as described above. Next, the OD_600_ was adjusted to 0.1 and 10 µl of culture was pipetted onto sterile polycarbonate membranes (25mm diameter, 0.2 µm pore size, MilliporeSigma™) on BHIYE-HP plates. Three polycarbonate membranes were placed on each plate, and five 10 µl spots of culture were pipetted onto each membrane for a total of 15 spots per plate. Two plates were prepared. One plate was incubated at 37 °C, 5% CO_2_ (for genes essential for colony growth in aerobic conditions), and the other was incubated in a Coy anaerobic chamber with an atmosphere of 90% N_2_, 5% CO_2_ and 5% H_2_ (for genes essential for colony growth in anaerobic conditions). After 24h of incubation, membranes were transferred to an Eppendorf tube and washed with sterile BHIYE medium. Cells were centrifuged (1min at 14,000g) and the pellets were collected for DNA extraction.

### RNAseq analysis

RNAseq reads obtained from a previous study (Perera et al., 2021) from *H. parainfluenzae* ATCC 33392 grown using the colony biofilm model were aligned, mapped, and differentially expressed genes were determined as previously described (Perera et al., 2021), with the exception that we re-sequenced the genome of *Hp* 392 and used this now-closed genome and its RAST annotation (Aziz et al., 2008), generated under default settings, as the reference, to allow direct comparison between the RNAseq and TnSeq datasets.

### TnSeq Data analysis

We utilized a previously established pipeline (Narayanan et al., 2017) with some modifications. All scripts utilized are available on GitHub (https://github.com/mramsey01/TnSeq-analysis-1). Prior to genome alignment, the initial sequence data was trimmed and filtered with Trimmomatic (Bolger et al., 2014) using a sliding window of four nucleotides and removing sequences when -phred33 quality score < 20, and when sequences were shorter than 30 nucleotides. These quality-filtered sequences were then aligned to their respective genomes using Bowtie2 (Langmead and Salzberg, 2012) under default settings. Next, *mariner* insertion sites and their associated read counts were calculated (Table S7). Local smoothing (LOESS) was used as previously described by (Turner et al., 2015) to correct for potential multifork chromosomal replication bias of insertion frequency. The Monte Carlo method was then applied as described previously (Lewin et al., 2019; Narayanan et al., 2017) to generate a pseudo dataset with expected number of insertions per TA sites. This method generates 100 pseudo datasets through sampling with replacement from the observed locations of the insertions and their respective read counts. Pseudo datasets were then compared to the observed distribution of read counts to determine where the number of observed read counts differs from the expected number of reads in the pseudo datasets. DESeq (Anders and Huber, 2010; Love et al., 2014) was used to compare the observed vs. expected (pseudo) datasets and calculate differential mutant abundance and significance. Then, following the protocols established by (Turner et al., 2015), a clustering algorithm was used to fit a bimodal curve on the distribution of calculated differential mutant abundances to cluster genes into two groups. Genes with substantially fewer reads were clustered under the “reduced” group, while genes for which the number of reads did not differ substantively from the expected number of reads were clustered under the “unchanged” group. The clustering algorithm also estimates an uncertainty measure for the assignment to each group. A gene was considered essential when three conditions were met: (A) the observed number of insertions was significantly (p-value <= 0.01) different from the expected number of insertions in the pseudo datasets; (B) the observed number of insertions was substantially smaller than the expected number of reads so that the gene was clustered under the “reduced” group, and; (C) the uncertainty of the placement of that gene in the reduced group was smaller than 0.01 (on a 0 to 1 scale).

## Discussion

Transposon sequencing (TnSeq) is a tool widely used to identify genes essential for survival (Barquist et al., 2013; Cain et al., 2020), many of which fall in the realm of “housekeeping” genes. This data alone has utility as a tool to identify potential drug targets (Mobegi et al., 2014). Further, TnSeq can be utilized in mutant strain backgrounds to find synthetically lethal genes involved in different synthesis or degradation pathways (Greene et al., 2018). TnSeq is also often used to compare different growth environments, e.g., *in vitro* vs. *in vivo*, finding conditionally essential genes, which are useful to generate hypothetical mechanisms for further testing and potential therapeutic targets (Turner et al., 2014).

Here, we use TnSeq to describe the essential genome and the conditionally essential genes necessary for *in vitro* biofilm growth under aerobic and anaerobic conditions in two strains of *Hp* (*Hp* 392 and *Hp* EL1). We chose to assess essential genomes using growth on a nutrient-rich medium so that accessory synthesis pathways would not be required and, thus, not deemed essential, as they would have been using a minimal, defined medium. This strategy allows us to identify only the most essential elements necessary for survival and allows for greater resolution of the completed library to interrogate other environments to find conditionally essential genes, making this a useful tool for future investigations. As expected, we observed that the majority of essential *Hp* genes are associated with central metabolic pathways and cell machinery, similar to results published in other studies (Lewin et al., 2019; Narayanan et al., 2017). We also identified a small subset of essential genes that were strain-specific (Table S7). Studies have demonstrated great genetic variation among *Hp* isolates. Thus, strain variation was an expected outcome and a motivating reason for performing these experiments (Kerr et al., 1993; Watts et al., 2021) with more than one strain. *Hp* 392 and *Hp* EL1 were isolated at very different times, geographic sites, and from distinct body sites. *Hp* 392 was isolated from a septic finger infection in London, England, around 1949, whereas *Hp* EL1 was isolated by our lab from the supragingival plaque of a healthy adult in Rhode Island in 2018. As the type-strain, *Hp* 392 was procured from the ATCC collection, and it is unknown how often it has been passaged prior to deposition and if the strain has been “domesticated” to an *in vitro* environment. The diverging histories of our two *Hp* strains ensures a greater likelihood that our data is more representative of *Hp* as a species. To further investigate the possibility of strain variation, we compared the genomes of our two strains here to 16 other *Hp* genomes available from NCBI (Fig 3). We found that all 341 essential genes shared between our strains were also present in each of the other genomes. This finding is similar to findings in *A. actinomycetemcomitans* (Narayanan et al., 2017) and indicates that the essential genes we identified are indeed part of the core of all the examined *Hp* genomes.

We would expect biofilm growth to require gene functions in addition to the absolute essential genes initially identified by TnSeq. We, therefore, used our libraries to identify conditionally essential genes for biofilm growth in aerobic and anaerobic conditions using a colony biofilm model (Anderl et al., 2000) grown on the same nutrient-rich medium that was used for the selection of the initial library. Most of the genes identified as essential for biofilm growth were essential regardless of oxygen availability. It is well-documented that in dense biofilms, oxygen gradients are formed through bacterial respiration (Ahn and Burne, 2007; Klementiev et al., 2020). While finding mutants that were conditionally essential for both oxygen conditions was expected, they were even more similar than we initially anticipated.

Some of the genes conditionally essential anaerobically for both *Hp* strains are the Na+-translocating NADH: quinone oxidoreductase (NQR) subunits A-F(*nqr*ABCDEF). NQR is a redox-driven sodium pump, well characterized in *Vibrio cholerae*, and operates in the respiratory chain, oxidizing cytoplasmic NADH and reducing it to ubiquinone (Agarwal et al., 2020; Steuber et al., 2014). Also anaerobically essential, were the four genes organized in operon that code for the fumarate reductase enzyme. It is unknown if this is acting as part of the reductive TCA cycle or if fumarate is serving as a terminal electron acceptor in this context (Kröger, 1978) and requires further investigation.

We were curious if the conditionally essential genes for biofilm growth would also be upregulated in biofilm growth. To assess this, we used our earlier transcriptome data (Perera et al., 2022) of colony biofilms under aerobic and anaerobic conditions for *Hp* 392 and compared genes differentially expressed in anaerobic biofilm growth to conditionally essential genes for anaerobic biofilm growth (Fig. 5). Interestingly, only three genes essential for anaerobic biofilm growth were significantly upregulated during anaerobiosis. Of those, two genes (Murein DD-endopeptidase and LPS-core synthesis glycosyltransferase) products are involved in cell metabolism, and one (Bacterial ribosome SSU maturation protein RimP) is involved in genetic information processing. The low overlap between gene essentiality and differential expression indicates that these conditionally essential genes are transcribed under aerobic conditions, where they are not essential, as well as anaerobic conditions, where they are essential. These data are similar to previous findings in *Pseudomonas aeruginosa* (Turner et al., 2014), indicating that mutant fitness and elevated gene expression are not well correlated. Likewise, other researchers have found that for *Listeria monocytogenes* (Müller-Herbst et al., 2014), genes significantly upregulated in anaerobiosis were not essential for anaerobic growth. It is also possible that basal level expression of many of these genes is sufficient for fitness in the short timeframes of our experimental model, but that regulation may be a factor under longer durations, where minor differences in efficiency are more relevant. Based on these results, we can argue that using transcriptomes to predict gene essentiality is not very effective. Increasing transcription of some genes may, however, contribute to how *Hp* adapts to changes in oxygen levels, allowing *Hp* to thrive in different portions of the biofilm.

In summary, we present the essential genomes for the type-strain and oral isolate of *H. parainfluenzae*, revealing that this essential genome is highly conserved between *Hp* strains. We also identified genes necessary for aerobic and anaerobic biofilm growth *in vitro*, showing that many genes involved in anaerobiosis are also essential for aerobic biofilm growth. Between these data and transcriptomes of aerobic vs. anaerobic growth, we determined that *Hp* grows in a highly heterogeneous environment for oxygen availability. These data may serve as a starting point for understanding *Hp* growth in oral biofilms and how it interacts with other members of polymicrobial oral biofilms in diverse niches within the oral cavity it inhabits.

## Acknowledgements

We would like to thank members of the M. Ramsey lab for valuable help with all phases of this manuscript. We thank Dr. Janet Atoyan in the URI Genomics Core Facility for assistance with sequencing library synthesis and quality control work.

