## Supplemental Figures and Tables for "Essential genes for *Haemophilus parainfluenzae* survival and biofilm growth"

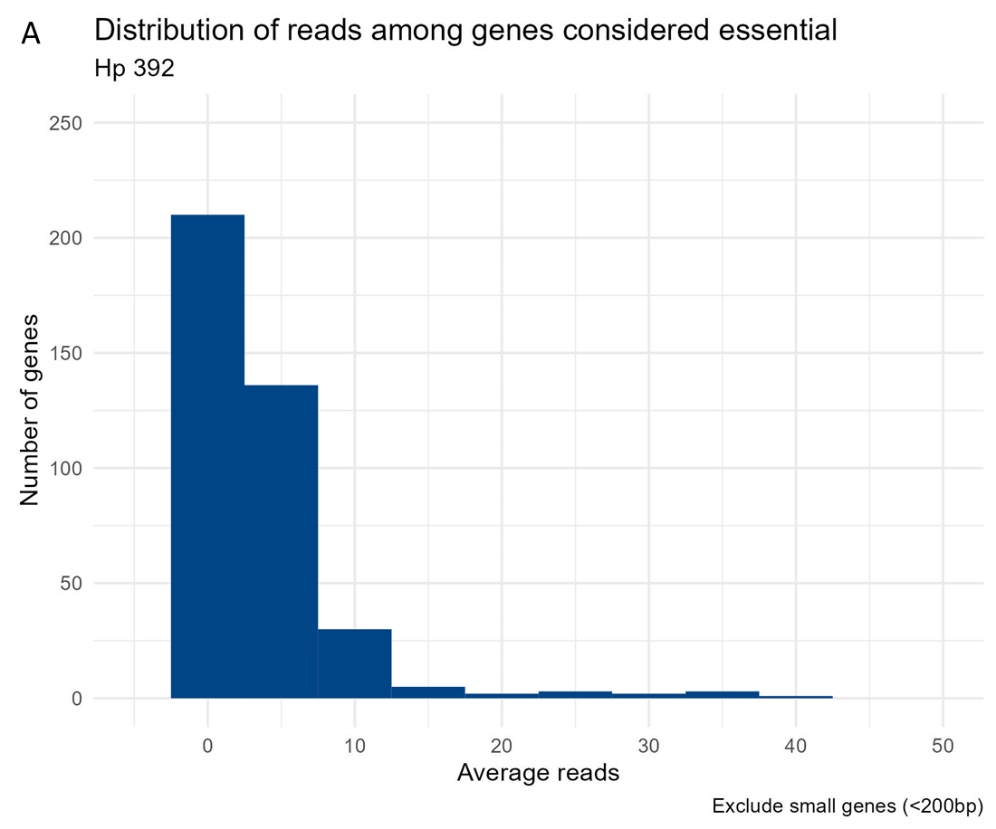

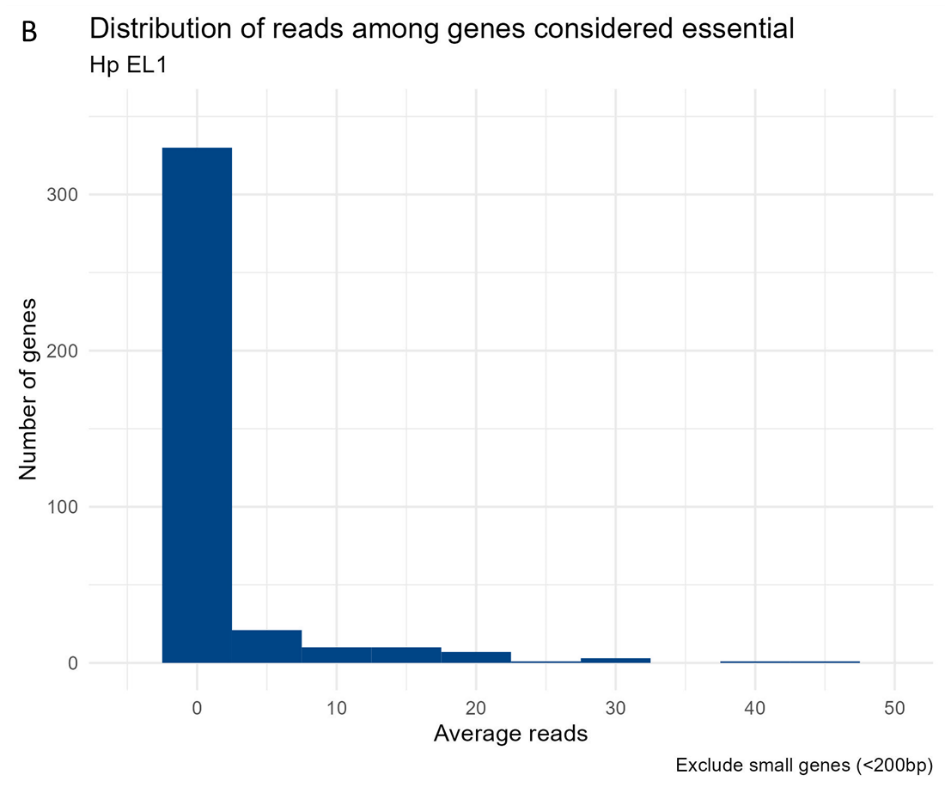

**Figure S1. Distribution of reads among genes considered essential *in vitro* for *Hp* 392 (A) and *Hp* EL1 (B).** Genes classified as essential had zero or significantly fewer insertions than non-essential genes.

**Table S1.** Strains and plasmids used.

| **Species** | **Identifier** | **Strain** | **Details** |
| --- | --- | --- | --- |
| *E. coli* | - | NEB5-ɑ | Cloning strain (New England Biolabs) |
| *E. coli* | MR0142 | MFD-pir | Donor strain Diaminopimelic acid auxotroph (Ferrières et al., 2010) |
| *H. parainfluenzae* | MR160 | ATCC33392 | *Haemophilus parainfluenzae* (ATCC 33392TM) |
| *H. parainfluenzae* | MR305 | EL1 | *Haemophilus parainfluenzae* isolated from healthy supragingival plaque |
| *H. parainfluenzae* | MR469 | ATCC33392 | Nalidixic acid resistant strain derived from ATCC33392 (this paper) |
| *H. parainfluenzae* | MR479 | EL1 | Nalidixic acid resistant strain derived from EL1 (this paper) |
| **Species** | **Identifier** | **plasmid** | **Details** |
| *E.coli* | MR042 | pMR361-K | Mariner delivery vector |

**Table S2.** List of primers used.

| **Identifier** | **Sequence (5’ – 3’)** | **Concentration** | **Barcode** | **Detail** |
| --- | --- | --- | --- | --- |
| MR163 | CTTTCAGACCGGGGACTTATCAGCCAACCTGTTA | 100um | - | Custom sequencing primer |
| MR165 | ACTCACTATAGGAGGGCGGGAATCATTTGAAGGTTGGTAC | 30uM | - | Mariner-specific PCR1 primer (5' biotin labelled) |
| MR164 | GTGACTGGAGTTCAGACGTGTGCTCTTCCGATCTGGGGGGGGGGGGGGGG | 30uM | - | Poly-C PCR1 primer |
| MR166 | AATGATACGGCGACCACCGAGATCTACACTCTTTcagaccggggacttatcagcc | 7.5uM | - | Mariner-specific PCR2, pair with any of barcode index primers below |
| MR167 | CAAGCAGAAGACGGCATACGAGATTCGCCTTAGTGACTGGAGTTCAGACGTGTGCTCTTCCGATCT | 7.5uM | BC73  (N701) | Barcode primer PCR2 |
| MR168 | CAAGCAGAAGACGGCATACGAGATTTCTGCCTGTGACTGGAGTTCAGACGTGTGCTCTTCCGATCT | 7.5uM | BC77  (N703) | Barcode primer PCR2 |
| MR169 | CAAGCAGAAGACGGCATACGAGATGCTCAGGAGTGACTGGAGTTCAGACGTGTGCTCTTCCGATCT | 7.5uM | BC79  (N704) | Barcode primer PCR2 |
| MR170 | CAAGCAGAAGACGGCATACGAGATAGGAGTCCGTGACTGGAGTTCAGACGTGTGCTCTTCCGATCT | 7.5uM | BC81  (N705) | Barcode primer PCR2 |
| MR171 | CAAGCAGAAGACGGCATACGAGATCATGCCTAGTGACTGGAGTTCAGACGTGTGCTCTTCCGATCT | 7.5uM | BC83  (N706) | Barcode primer PCR2 |
| MR172 | CAAGCAGAAGACGGCATACGAGATGTAGAGAGGTGACTGGAGTTCAGACGTGTGCTCTTCCGATCT | 7.5uM | BC85  (N707) | Barcode primer PCR2 |
| MR173 | CAAGCAGAAGACGGCATACGAGATCCTCTCTGGTGACTGGAGTTCAGACGTGTGCTCTTCCGATCT | 7.5uM | BC87  (N708) | Barcode primer PCR2 |
| MR174 | CAAGCAGAAGACGGCATACGAGATAGCGTAGCGTGACTGGAGTTCAGACGTGTGCTCTTCCGATCT | 7.5uM | BC89  (N709) | Barcode primer PCR2 |
| MR175 | CAAGCAGAAGACGGCATACGAGATCAGCCTCGGTGACTGGAGTTCAGACGTGTGCTCTTCCGATCT | 7.5uM | BC90  (N710) | Barcode primer PCR2 |
| MR176 | CAAGCAGAAGACGGCATACGAGATTGCCTCTTGTGACTGGAGTTCAGACGTGTGCTCTTCCGATCT | 7.5uM | BC91  (N711) | Barcode primer PCR2 |
| MR653 | CAAGCAGAAGACGGCATACGAGATCCTGAGATGTGACTGGAGTTCAGACGTGTGCTCTTCCGATCT | 7.5uM | H/N 712 | Barcode primer PCR2 |
| MR654 | CAAGCAGAAGACGGCATACGAGATCCTGAGATGTGACTGGAGTTCAGACGTGTGCTCTTCCGATCT | 7.5uM | H/N 715 | Barcode primer PCR2 |
| MR655 | CAAGCAGAAGACGGCATACGAGATTAGCGAGTGTGACTGGAGTTCAGACGTGTGCTCTTCCGATCT | 7.5uM | H/N 716 | Barcode primer PCR2 |
| MR656 | CAAGCAGAAGACGGCATACGAGATTACTACGCGTGACTGGAGTTCAGACGTGTGCTCTTCCGATCT | 7.5uM | H/N 719 | Barcode primer PCR2 |
| MR657 | CAAGCAGAAGACGGCATACGAGATAGGCTCCGGTGACTGGAGTTCAGACGTGTGCTCTTCCGATCT | 7.5uM | H/N 720 | Barcode primer PCR2 |

**Table S3.** Anaerobic conditionally essential genes in *Hp* 392 and their predicted KEGG functions.

| **PEG** | **Annotation** | **KEGG lowest category** | **KEGG highest category** |
| --- | --- | --- | --- |
| 1681 | Glucose-6-phosphate isomerase (EC 5.3.1.9) | Glycolysis / Gluconeogenesis | Carbohydrate metabolism |
| 373 | Phosphoenolpyruvate carboxykinase [ATP] (EC 4.1.1.49) | Glycolysis / Gluconeogenesis | Carbohydrate metabolism |
| 1940 | PTS system glucose-specific IIA component (EC 2.7.1.199) | Glycolysis / Gluconeogenesis | Carbohydrate metabolism |
| 908 | Fumarate reductase flavoprotein subunit (EC 1.3.5.4) | Citrate cycle (TCA cycle) | Carbohydrate metabolism |
| 907 | Fumarate reductase iron-sulfur protein (EC 1.3.5.4) | Citrate cycle (TCA cycle) | Carbohydrate metabolism |
| 906 | Fumarate reductase subunit C | Citrate cycle (TCA cycle) | Carbohydrate metabolism |
| 905 | Fumarate reductase subunit D | Citrate cycle (TCA cycle) | Carbohydrate metabolism |
| 1934 | Fumarate hydratase class II (EC 4.2.1.2) | Citrate cycle (TCA cycle) | Carbohydrate metabolism |
| 1788 | Malate dehydrogenase (EC 1.1.1.37) | Citrate cycle (TCA cycle) | Carbohydrate metabolism |
| 373 | Phosphoenolpyruvate carboxykinase [ATP] (EC 4.1.1.49) | Citrate cycle (TCA cycle) | Carbohydrate metabolism |
| 1681 | Glucose-6-phosphate isomerase (EC 5.3.1.9) | Pentose phosphate pathway | Carbohydrate metabolism |
| 145 | Transketolase (EC 2.2.1.1) | Pentose phosphate pathway | Carbohydrate metabolism |
| 502 | Ribose-5-phosphate isomerase A (EC 5.3.1.6) | Pentose phosphate pathway | Carbohydrate metabolism |
| 1940 | PTS system glucose-specific IIA component (EC 2.7.1.199) | Starch and sucrose metabolism | Carbohydrate metabolism |
| 1681 | Glucose-6-phosphate isomerase (EC 5.3.1.9) | Starch and sucrose metabolism | Carbohydrate metabolism |
| 1940 | PTS system glucose-specific IIA component (EC 2.7.1.199) | Amino sugar and nucleotide sugar metabolism | Carbohydrate metabolism |
| 4 | N-acetylglucosamine-6-phosphate deacetylase (EC 3.5.1.25) | Amino sugar and nucleotide sugar metabolism | Carbohydrate metabolism |
| 1681 | Glucose-6-phosphate isomerase (EC 5.3.1.9) | Amino sugar and nucleotide sugar metabolism | Carbohydrate metabolism |
| 292 | Pyruvate formate-lyase (EC 2.3.1.54) | Pyruvate metabolism | Carbohydrate metabolism |
| 1788 | Malate dehydrogenase (EC 1.1.1.37) | Pyruvate metabolism | Carbohydrate metabolism |
| 1934 | Fumarate hydratase class II (EC 4.2.1.2) | Pyruvate metabolism | Carbohydrate metabolism |
| 908 | Fumarate reductase flavoprotein subunit (EC 1.3.5.4) | Pyruvate metabolism | Carbohydrate metabolism |
| 907 | Fumarate reductase iron-sulfur protein (EC 1.3.5.4) | Pyruvate metabolism | Carbohydrate metabolism |
| 906 | Fumarate reductase subunit C | Pyruvate metabolism | Carbohydrate metabolism |
| 905 | Fumarate reductase subunit D | Pyruvate metabolism | Carbohydrate metabolism |
| 373 | Phosphoenolpyruvate carboxykinase [ATP] (EC 4.1.1.49) | Pyruvate metabolism | Carbohydrate metabolism |
| 1788 | Malate dehydrogenase (EC 1.1.1.37) | Glyoxylate and dicarboxylate metabolism | Carbohydrate metabolism |
| 292 | Pyruvate formate-lyase (EC 2.3.1.54) | Propanoate metabolism | Carbohydrate metabolism |
| 908 | Fumarate reductase flavoprotein subunit (EC 1.3.5.4) | Butanoate metabolism | Carbohydrate metabolism |
| 907 | Fumarate reductase iron-sulfur protein (EC 1.3.5.4) | Butanoate metabolism | Carbohydrate metabolism |
| 906 | Fumarate reductase subunit C | Butanoate metabolism | Carbohydrate metabolism |
| 905 | Fumarate reductase subunit D | Butanoate metabolism | Carbohydrate metabolism |
| 292 | Pyruvate formate-lyase (EC 2.3.1.54) | Butanoate metabolism | Carbohydrate metabolism |
| 908 | Fumarate reductase flavoprotein subunit (EC 1.3.5.4) | Oxidative phosphorylation | Energy metabolism |
| 907 | Fumarate reductase iron-sulfur protein (EC 1.3.5.4) | Oxidative phosphorylation | Energy metabolism |
| 906 | Fumarate reductase subunit C | Oxidative phosphorylation | Energy metabolism |
| 905 | Fumarate reductase subunit D | Oxidative phosphorylation | Energy metabolism |
| 575 | ATP synthase alpha chain (EC 3.6.3.14) | Oxidative phosphorylation | Energy metabolism |
| 577 | ATP synthase beta chain (EC 3.6.3.14) | Oxidative phosphorylation | Energy metabolism |
| 576 | ATP synthase gamma chain (EC 3.6.3.14) | Oxidative phosphorylation | Energy metabolism |
| 574 | ATP synthase delta chain (EC 3.6.3.14) | Oxidative phosphorylation | Energy metabolism |
| 571 | ATP synthase F0 sector subunit a (EC 3.6.3.14) | Oxidative phosphorylation | Energy metabolism |
| 573 | ATP synthase F0 sector subunit b (EC 3.6.3.14) | Oxidative phosphorylation | Energy metabolism |
| 572 | ATP synthase F0 sector subunit c (EC 3.6.3.14) | Oxidative phosphorylation | Energy metabolism |
| 577 | ATP synthase beta chain (EC 3.6.3.14) | Photosynthesis | Energy metabolism |
| 576 | ATP synthase gamma chain (EC 3.6.3.14) | Photosynthesis | Energy metabolism |
| 575 | ATP synthase alpha chain (EC 3.6.3.14) | Photosynthesis | Energy metabolism |
| 574 | ATP synthase delta chain (EC 3.6.3.14) | Photosynthesis | Energy metabolism |
| 573 | ATP synthase F0 sector subunit b (EC 3.6.3.14) | Photosynthesis | Energy metabolism |
| 572 | ATP synthase F0 sector subunit c (EC 3.6.3.14) | Photosynthesis | Energy metabolism |
| 571 | ATP synthase F0 sector subunit a (EC 3.6.3.14) | Photosynthesis | Energy metabolism |
| 145 | Transketolase (EC 2.2.1.1) | Carbon fixation in photosynthetic organisms | Energy metabolism |
| 502 | Ribose-5-phosphate isomerase A (EC 5.3.1.6) | Carbon fixation in photosynthetic organisms | Energy metabolism |
| 373 | Phosphoenolpyruvate carboxykinase [ATP] (EC 4.1.1.49) | Carbon fixation in photosynthetic organisms | Energy metabolism |
| 1788 | Malate dehydrogenase (EC 1.1.1.37) | Carbon fixation in photosynthetic organisms | Energy metabolism |
| 1788 | Malate dehydrogenase (EC 1.1.1.37) | Carbon fixation pathways in prokaryotes | Energy metabolism |
| 1934 | Fumarate hydratase class II (EC 4.2.1.2) | Carbon fixation pathways in prokaryotes | Energy metabolism |
| 908 | Fumarate reductase flavoprotein subunit (EC 1.3.5.4) | Carbon fixation pathways in prokaryotes | Energy metabolism |
| 907 | Fumarate reductase iron-sulfur protein (EC 1.3.5.4) | Carbon fixation pathways in prokaryotes | Energy metabolism |
| 906 | Fumarate reductase subunit C | Carbon fixation pathways in prokaryotes | Energy metabolism |
| 905 | Fumarate reductase subunit D | Carbon fixation pathways in prokaryotes | Energy metabolism |
| 1788 | Malate dehydrogenase (EC 1.1.1.37) | Methane metabolism | Energy metabolism |
| 1315 | Deoxycytidine triphosphate deaminase (EC 3.5.4.13) | Pyrimidine metabolism | Nucleotide metabolism |
| 1788 | Malate dehydrogenase (EC 1.1.1.37) | Cysteine and methionine metabolism | Amino acid metabolism |
| 802 | D-glycero-beta-D-manno-heptose-7-bisphosphate 7-phosphatase | Lipopolysaccharide biosynthesis | Glycan biosynthesis and metabolism |
| 1330 | Biotin operon repressor / Biotin--protein ligase | Biotin metabolism | Metabolism of cofactors and vitamins |
| 32 | 2-succinyl-6-hydroxy-cyclohexadiene-1-carboxylate synthase | Ubiquinone and other terpenoid-quinone biosynthesis | Metabolism of cofactors and vitamins |
| 1854 | O-succinylbenzoic acid--CoA ligase (EC 6.2.1.26) | Ubiquinone and other terpenoid-quinone biosynthesis | Metabolism of cofactors and vitamins |
| 145 | Transketolase (EC 2.2.1.1) | Biosynthesis of ansamycins | Metabolism of terpenoids and polyketides |
| 1814 | SSU ribosomal protein S15p (S13e) | Ribosome | Translation |
| 1166 | Arginine ABC transporter%2C substrate-binding protein ArtI | ABC transporters | Membrane transport |
| 1940 | PTS system glucose-specific IIA component (EC 2.7.1.199) | Phosphotransferase system (PTS) | Membrane transport |
| 1238 | PTS IIA-like nitrogen-regulatory protein PtsN | Phosphotransferase system (PTS) | Membrane transport |
| 908 | Fumarate reductase flavoprotein subunit (EC 1.3.5.4) | Two-component system | Signal transduction |
| 907 | Fumarate reductase iron-sulfur protein (EC 1.3.5.4) | Two-component system | Signal transduction |
| 906 | Fumarate reductase subunit C | Two-component system | Signal transduction |
| 905 | Fumarate reductase subunit D | Two-component system | Signal transduction |
| 93 | Nitrogen regulatory protein P-II | Two-component system | Signal transduction |
| 11 | Cyclic AMP receptor protein | Two-component system | Signal transduction |
| 11 | Cyclic AMP receptor protein | Quorum sensing | Cellular community - prokaryotes |
| 1940 | PTS system glucose-specific IIA component (EC 2.7.1.199) | Biofilm formation - Vibrio cholerae | Cellular community - prokaryotes |
| 11 | Cyclic AMP receptor protein | Biofilm formation - Vibrio cholerae | Cellular community - prokaryotes |
| 11 | Cyclic AMP receptor protein | Biofilm formation - Pseudomonas aeruginosa | Cellular community - prokaryotes |
| 1940 | PTS system glucose-specific IIA component (EC 2.7.1.199) | Biofilm formation - Escherichia coli | Cellular community - prokaryotes |
| 11 | Cyclic AMP receptor protein | Biofilm formation - Escherichia coli | Cellular community - prokaryotes |
| 1278 | RNA polymerase-binding transcription factor DksA | Biofilm formation - Escherichia coli | Cellular community - prokaryotes |
| 1934 | Fumarate hydratase class II (EC 4.2.1.2) | Pathways in cancer | Other |
| 1934 | Fumarate hydratase class II (EC 4.2.1.2) | Renal cell carcinoma | Other |
| 1934 | Fumarate hydratase class II (EC 4.2.1.2) | Cushing syndrome | Other |
| 1030 | Murein DD-endopeptidase MepM | Peptidases and inhibitors [BR:ko | Protein families: metabolism |
| 802 | D-glycero-beta-D-manno-heptose-1%2C7-bisphosphate 7-phosphatase | Lipopolysaccharide biosynthesis proteins [BR:ko | Protein families: metabolism |
| 1030 | Murein DD-endopeptidase MepM | Peptidoglycan biosynthesis and degradation proteins | Protein families: metabolism |
| 577 | ATP synthase beta chain (EC 3.6.3.14) | Photosynthesis proteins [BR:ko | Protein families: metabolism |
| 576 | ATP synthase gamma chain (EC 3.6.3.14) | Photosynthesis proteins [BR:ko | Protein families: metabolism |
| 575 | ATP synthase alpha chain (EC 3.6.3.14) | Photosynthesis proteins [BR:ko | Protein families: metabolism |
| 574 | ATP synthase delta chain (EC 3.6.3.14) | Photosynthesis proteins [BR:ko | Protein families: metabolism |
| 573 | ATP synthase F0 sector subunit b (EC 3.6.3.14) | Photosynthesis proteins [BR:ko | Protein families: metabolism |
| 572 | ATP synthase F0 sector subunit c (EC 3.6.3.14) | Photosynthesis proteins [BR:ko | Protein families: metabolism |
| 571 | ATP synthase F0 sector subunit a (EC 3.6.3.14) | Photosynthesis proteins [BR:ko | Protein families: metabolism |
| 570 | Permease of the drug/metabolite transporter (DMT) superfamily | Photosynthesis proteins [BR:ko | Protein families: metabolism |
| 11 | Cyclic AMP receptor protein | Transcription factors [BR:ko | Protein families: genetic information processing |
| 1330 | Biotin operon repressor / Biotin--protein ligase | Transcription factors [BR:ko | Protein families: genetic information processing |
| 1278 | RNA polymerase-binding transcription factor DksA | Transcription factors [BR:ko | Protein families: genetic information processing |
| 1278 | RNA polymerase-binding transcription factor DksA | Transcription machinery [BR:ko | Protein families: genetic information processing |
| 1814 | SSU ribosomal protein S15p (S13e) | Ribosome [BR:ko | Protein families: genetic information processing |
| 1278 | RNA polymerase-binding transcription factor DksA | Ribosome biogenesis [BR:ko | Protein families: genetic information processing |
| 1625 | Bacterial ribosome SSU maturation protein RimP | Ribosome biogenesis [BR:ko | Protein families: genetic information processing |
| 1291 | Metal-dependent hydrolase YbeY | Ribosome biogenesis [BR:ko | Protein families: genetic information processing |
| 2002 | Ribonuclease P protein component (EC 3.1.26.5) | Transfer RNA biogenesis [BR:ko | Protein families: genetic information processing |
| 571 | ATP synthase F0 sector subunit a (EC 3.6.3.14) | Chaperones and folding catalysts [BR:ko | Protein families: genetic information processing |
| 2010 | Plasmid partitioning protein ParA | Chromosome and associated proteins [BR:ko | Protein families: genetic information processing |
| 1166 | Arginine ABC transporter substrate-binding protein ArtI | Transporters [BR:ko | Protein families: signaling and cellular processes |
| 1940 | PTS system%2C glucose-specific IIA component (EC 2.7.1.199) | Transporters [BR:ko | Protein families: signaling and cellular processes |
| 1238 | PTS IIA-like nitrogen-regulatory protein PtsN | Transporters [BR:ko | Protein families: signaling and cellular processes |
| 862 | Pantothenate:Na+ symporter (TC 2.A.21.1.1) | Transporters [BR:ko | Protein families: signaling and cellular processes |
| 924 | Serine transporter | Transporters [BR:ko | Protein families: signaling and cellular processes |
| 2010 | Plasmid partitioning protein ParA | Cytoskeleton proteins [BR:ko | Protein families: signaling and cellular processes |
| 1681 | Glucose-6-phosphate isomerase (EC 5.3.1.9) | Exosome [BR:ko | Protein families: signaling and cellular processes |
| 290 | Pyruvate formate-lyase activating enzyme (EC 1.97.1.4) | Enzymes with EC numbers | Unclassified: metabolism |
| 186 | Na(+)-translocating NADH-quinone reductase subunit A (EC 1.6.5.8) | Enzymes with EC numbers | Unclassified: metabolism |
| 185 | Na(+)-translocating NADH-quinone reductase subunit B (EC 1.6.5.8) | Enzymes with EC numbers | Unclassified: metabolism |
| 184 | Na(+)-translocating NADH-quinone reductase subunit C (EC 1.6.5.8) | Enzymes with EC numbers | Unclassified: metabolism |
| 183 | Na(+)-translocating NADH-quinone reductase subunit D (EC 1.6.5.8) | Enzymes with EC numbers | Unclassified: metabolism |
| 182 | Na(+)-translocating NADH-quinone reductase subunit E (EC 1.6.5.8) | Enzymes with EC numbers | Unclassified: metabolism |
| 181 | Na(+)-translocating NADH-quinone reductase subunit F (EC 1.6.5.8) | Enzymes with EC numbers | Unclassified: metabolism |
| 1703 | probable membrane protein YPO2224 | Glycan metabolism | Unclassified: metabolism |
| 1429 | LPS-core synthesis glycosyltransferase PM0509 | Glycan metabolism | Unclassified: metabolism |
| 659 | Murein hydrolase activator EnvC | Cell growth | Unclassified: signaling and cellular processes |
| 385 | Murein hydrolase activator NlpD | Structural proteins | Unclassified: signaling and cellular processes |
| 1374 | Penicillin-binding protein activator LpoA | Function unknown | Poorly characterized |
| 2001 | Membrane protein insertion efficiency factor YidD | Function unknown | Poorly characterized |
| 73 | Proposed lipoate regulatory protein YbeD | Function unknown | Poorly characterized |

**Table S4.** List of conditionally essential genes in *Hp* EL1 and their predicted KEGG functions.

| **PEG** | **Annotation** | **KEGG lowest category** | **KEGG highest category** |
| --- | --- | --- | --- |
| 1691 | Glucose-6-phosphate isomerase (EC 5.3.1.9) | Glycolysis / Gluconeogenesis | Carbohydrate metabolism |
| 1914 | Pyruvate dehydrogenase E1 component (EC 1.2.4.1) | Glycolysis / Gluconeogenesis | Carbohydrate metabolism |
| 1913 | Dihydrolipoamide acetyltransferase component of pyruvate dehydrogenase complex | Glycolysis / Gluconeogenesis | Carbohydrate metabolism |
| 221 | Glucose-1-phosphatase (EC 3.1.3.10) | Glycolysis / Gluconeogenesis | Carbohydrate metabolism |
| 1957 | PTS system glucose-specific IIA component (EC 2.7.1.199) | Glycolysis / Gluconeogenesis | Carbohydrate metabolism |
| 1679 | Dihydrolipoamide succinyltransferase component (E2) of 2-oxoglutarate dehydrogenase complex | Citrate cycle (TCA cycle) | Carbohydrate metabolism |
| 858 | Fumarate reductase flavoprotein subunit (EC 1.3.5.4) | Citrate cycle (TCA cycle) | Carbohydrate metabolism |
| 857 | Fumarate reductase iron-sulfur protein (EC 1.3.5.4) | Citrate cycle (TCA cycle) | Carbohydrate metabolism |
| 855 | Fumarate reductase subunit D | Citrate cycle (TCA cycle) | Carbohydrate metabolism |
| 1914 | Pyruvate dehydrogenase E1 component (EC 1.2.4.1) | Citrate cycle (TCA cycle) | Carbohydrate metabolism |
| 1913 | Dihydrolipoamide acetyltransferase component of pyruvate dehydrogenase complex | Citrate cycle (TCA cycle) | Carbohydrate metabolism |
| 1691 | Glucose-6-phosphate isomerase (EC 5.3.1.9) | Pentose phosphate pathway | Carbohydrate metabolism |
| 661 | Ribulose-phosphate 3-epimerase (EC 5.1.3.1) | Pentose phosphate pathway | Carbohydrate metabolism |
| 151 | Transketolase (EC 2.2.1.1) | Pentose phosphate pathway | Carbohydrate metabolism |
| 460 | Ribose-5-phosphate isomerase A (EC 5.3.1.6) | Pentose phosphate pathway | Carbohydrate metabolism |
| 661 | Ribulose-phosphate 3-epimerase (EC 5.1.3.1) | Pentose and glucuronate interconversions | Carbohydrate metabolism |
| 1957 | PTS system glucose-specific IIA component (EC 2.7.1.199) | Starch and sucrose metabolism | Carbohydrate metabolism |
| 1691 | Glucose-6-phosphate isomerase (EC 5.3.1.9) | Starch and sucrose metabolism | Carbohydrate metabolism |
| 1957 | PTS system glucose-specific IIA component (EC 2.7.1.199) | Amino sugar and nucleotide sugar metabolism | Carbohydrate metabolism |
| 1691 | Glucose-6-phosphate isomerase (EC 5.3.1.9) | Amino sugar and nucleotide sugar metabolism | Carbohydrate metabolism |
| 1914 | Pyruvate dehydrogenase E1 component (EC 1.2.4.1) | Pyruvate metabolism | Carbohydrate metabolism |
| 1913 | Dihydrolipoamide acetyltransferase component of pyruvate dehydrogenase complex | Pyruvate metabolism | Carbohydrate metabolism |
| 1428 | Acetate kinase (EC 2.7.2.1) | Pyruvate metabolism | Carbohydrate metabolism |
| 1427 | BioD-like N-terminal domain / Phosphate acetyltransferase (EC 2.3.1.8) | Pyruvate metabolism | Carbohydrate metabolism |
| 858 | Fumarate reductase flavoprotein subunit (EC 1.3.5.4) | Pyruvate metabolism | Carbohydrate metabolism |
| 857 | Fumarate reductase iron-sulfur protein (EC 1.3.5.4) | Pyruvate metabolism | Carbohydrate metabolism |
| 855 | Fumarate reductase subunit D | Pyruvate metabolism | Carbohydrate metabolism |
| 1428 | Acetate kinase (EC 2.7.2.1) | Propanoate metabolism | Carbohydrate metabolism |
| 1427 | BioD-like N-terminal domain / Phosphate acetyltransferase (EC 2.3.1.8) | Propanoate metabolism | Carbohydrate metabolism |
| 858 | Fumarate reductase flavoprotein subunit (EC 1.3.5.4) | Butanoate metabolism | Carbohydrate metabolism |
| 857 | Fumarate reductase iron-sulfur protein (EC 1.3.5.4) | Butanoate metabolism | Carbohydrate metabolism |
| 855 | Fumarate reductase subunit D | Butanoate metabolism | Carbohydrate metabolism |
| 858 | Fumarate reductase flavoprotein subunit (EC 1.3.5.4) | Oxidative phosphorylation | Energy metabolism |
| 857 | Fumarate reductase iron-sulfur protein (EC 1.3.5.4) | Oxidative phosphorylation | Energy metabolism |
| 855 | Fumarate reductase subunit D | Oxidative phosphorylation | Energy metabolism |
| 532 | ATP synthase alpha chain (EC 3.6.3.14) | Oxidative phosphorylation | Energy metabolism |
| 534 | ATP synthase beta chain (EC 3.6.3.14) | Oxidative phosphorylation | Energy metabolism |
| 533 | ATP synthase gamma chain (EC 3.6.3.14) | Oxidative phosphorylation | Energy metabolism |
| 531 | ATP synthase delta chain (EC 3.6.3.14) | Oxidative phosphorylation | Energy metabolism |
| 528 | ATP synthase F0 sector subunit a (EC 3.6.3.14) | Oxidative phosphorylation | Energy metabolism |
| 529 | ATP synthase F0 sector subunit c (EC 3.6.3.14) | Oxidative phosphorylation | Energy metabolism |
| 534 | ATP synthase beta chain (EC 3.6.3.14) | Photosynthesis | Energy metabolism |
| 533 | ATP synthase gamma chain (EC 3.6.3.14) | Photosynthesis | Energy metabolism |
| 532 | ATP synthase alpha chain (EC 3.6.3.14) | Photosynthesis | Energy metabolism |
| 531 | ATP synthase delta chain (EC 3.6.3.14) | Photosynthesis | Energy metabolism |
| 529 | ATP synthase F0 sector subunit c (EC 3.6.3.14) | Photosynthesis | Energy metabolism |
| 528 | ATP synthase F0 sector subunit a (EC 3.6.3.14) | Photosynthesis | Energy metabolism |
| 151 | Transketolase (EC 2.2.1.1) | Carbon fixation in photosynthetic organisms | Energy metabolism |
| 460 | Ribose-5-phosphate isomerase A (EC 5.3.1.6) | Carbon fixation in photosynthetic organisms | Energy metabolism |
| 661 | Ribulose-phosphate 3-epimerase (EC 5.1.3.1) | Carbon fixation in photosynthetic organisms | Energy metabolism |
| 858 | Fumarate reductase flavoprotein subunit (EC 1.3.5.4) | Carbon fixation pathways in prokaryotes | Energy metabolism |
| 857 | Fumarate reductase iron-sulfur protein (EC 1.3.5.4) | Carbon fixation pathways in prokaryotes | Energy metabolism |
| 855 | Fumarate reductase subunit D | Carbon fixation pathways in prokaryotes | Energy metabolism |
| 1427 | BioD-like N-terminal domain / Phosphate acetyltransferase (EC 2.3.1.8) | Carbon fixation pathways in prokaryotes | Energy metabolism |
| 1428 | Acetate kinase (EC 2.7.2.1) | Carbon fixation pathways in prokaryotes | Energy metabolism |
| 1428 | Acetate kinase (EC 2.7.2.1) | Methane metabolism | Energy metabolism |
| 1427 | BioD-like N-terminal domain / Phosphate acetyltransferase (EC 2.3.1.8) | Methane metabolism | Energy metabolism |
| 1443 | Carbonic anhydrase%2C beta class (EC 4.2.1.1) | Nitrogen metabolism | Energy metabolism |
| 477 | Serine acetyltransferase (EC 2.3.1.30) | Sulfur metabolism | Energy metabolism |
| 1356 | Cardiolipin synthase bacterial type ClsA | Glycerophospholipid metabolism | Lipid metabolism |
| 172 | 5'-nucleotidase (EC 3.1.3.5)%3B NAD pyrophosphatase periplasmic (EC 3.6.1.22) | Purine metabolism | Nucleotide metabolism |
| 172 | 5'-nucleotidase (EC 3.1.3.5)%3B NAD pyrophosphatase periplasmic (EC 3.6.1.22) | Pyrimidine metabolism | Nucleotide metabolism |
| 1047 | Threonine synthase (EC 4.2.3.1) | Glycine, serine and threonine metabolism | Amino acid metabolism |
| 477 | Serine acetyltransferase (EC 2.3.1.30) | Cysteine and methionine metabolism | Amino acid metabolism |
| 1679 | Dihydrolipoamide succinyltransferase component (E2) of 2-oxoglutarate dehydrogenase complex | Lysine degradation | Amino acid metabolism |
| 1679 | Dihydrolipoamide succinyltransferase component (E2) of 2-oxoglutarate dehydrogenase complex | Tryptophan metabolism | Amino acid metabolism |
| 1365 | 2-keto-3-deoxy-D-arabino-heptulosonate-7-phosphate synthase I alpha (EC 2.5.1.54) | Phenylalanine, tyrosine and tryptophan biosynthesis | Amino acid metabolism |
| 626 | Shikimate 5-dehydrogenase I alpha (EC 1.1.1.25) | Phenylalanine, tyrosine and tryptophan biosynthesis | Amino acid metabolism |
| 1352 | 3-phosphoshikimate 1-carboxyvinyltransferase (EC 2.5.1.19) | Phenylalanine, tyrosine and tryptophan biosynthesis | Amino acid metabolism |
| 1875 | Chorismate synthase (EC 4.2.3.5) | Phenylalanine, tyrosine and tryptophan biosynthesis | Amino acid metabolism |
| 1427 | BioD-like N-terminal domain / Phosphate acetyltransferase (EC 2.3.1.8) | Taurine and hypotaurine metabolism | Metabolism of other amino acids |
| 1428 | Acetate kinase (EC 2.7.2.1) | Taurine and hypotaurine metabolism | Metabolism of other amino acids |
| 932 | Acyl-[acyl-carrier-protein]--UDP-N-acetylglucosamine O-acyltransferase (EC 2.3.1.129) | Lipopolysaccharide biosynthesis | Glycan biosynthesis and metabolism |
| 1783 | D-arabinose-5-phosphate isomerase (EC 5.3.1.13) | Lipopolysaccharide biosynthesis | Glycan biosynthesis and metabolism |
| 1128 | D-sedoheptulose 7-phosphate isomerase (EC 5.3.1.28) | Lipopolysaccharide biosynthesis | Glycan biosynthesis and metabolism |
| 1851 | D-glycero-beta-D-manno-heptose 1-phosphate adenylyltransferase | Lipopolysaccharide biosynthesis | Glycan biosynthesis and metabolism |
| 339 | ADP-heptose--lipooligosaccharide heptosyltransferase II | Lipopolysaccharide biosynthesis | Glycan biosynthesis and metabolism |
| 1155 | UDP-N-acetylglucosamine--N-acetylmuramyl-(pentapeptide) | Peptidoglycan biosynthesis | Glycan biosynthesis and metabolism |
| 1149 | Cell division protein FtsI [Peptidoglycan synthetase] (EC 2.4.1.129) | Peptidoglycan biosynthesis | Glycan biosynthesis and metabolism |
| 1592 | Thiamine-monophosphate kinase (EC 2.7.4.16) | Thiamine metabolism | Metabolism of cofactors and vitamins |
| 639 | 3%2C4-dihydroxy-2-butanone 4-phosphate synthase (EC 4.1.99.12) | Riboflavin metabolism | Metabolism of cofactors and vitamins |
| 1047 | Threonine synthase (EC 4.2.3.1) | Vitamin B | Metabolism of cofactors and vitamins |
| 172 | 5'-nucleotidase (EC 3.1.3.5)%3B NAD pyrophosphatase periplasmic (EC 3.6.1.22) | Nicotinate and nicotinamide metabolism | Metabolism of cofactors and vitamins |
| 73 | Octanoate-[acyl-carrier-protein]-protein-N-octanoyltransferase (EC 2.3.1.181) | Lipoic acid metabolism | Metabolism of cofactors and vitamins |
| 1420 | Glutamate-1-semialdehyde 2%2C1-aminomutase (EC 5.4.3.8) | Porphyrin and chlorophyll metabolism | Metabolism of cofactors and vitamins |
| 861 | Ferrochelatase%2C protoheme ferro-lyase (EC 4.99.1.1) | Porphyrin and chlorophyll metabolism | Metabolism of cofactors and vitamins |
| 32 | 2-succinyl-6-hydroxy-4-cyclohexadiene-1-carboxylate synthase (EC 4.2.99.20) | Ubiquinone and other terpenoid-quinone biosynthesis | Metabolism of cofactors and vitamins |
| 805 | O-succinylbenzoate synthase (EC 4.2.1.113) | Ubiquinone and other terpenoid-quinone biosynthesis | Metabolism of cofactors and vitamins |
| 1877 | O-succinylbenzoic acid--CoA ligase (EC 6.2.1.26) | Ubiquinone and other terpenoid-quinone biosynthesis | Metabolism of cofactors and vitamins |
| 804 | Naphthoate synthase (EC 4.1.3.36) | Ubiquinone and other terpenoid-quinone biosynthesis | Metabolism of cofactors and vitamins |
| 862 | 1%2C4-dihydroxy-2-naphthoyl-CoA hydrolase (EC 3.1.2.28) in menaquinone biosynthesis | Ubiquinone and other terpenoid-quinone biosynthesis | Metabolism of cofactors and vitamins |
| 269 | 4-hydroxy-3-methylbut-2-enyl diphosphate reductase (EC 1.17.7.4) | Terpenoid backbone biosynthesis | Metabolism of terpenoids and polyketides |
| 1965 | tRNA dimethylallyltransferase (EC 2.5.1.75) | Zeatin biosynthesis | Metabolism of terpenoids and polyketides |
| 151 | Transketolase (EC 2.2.1.1) | Biosynthesis of ansamycins | Metabolism of terpenoids and polyketides |
| 385 | LSU ribosomal protein L17p | Ribosome | Translation |
| 1602 | Glutaminyl-tRNA synthetase (EC 6.1.1.18) | Aminoacyl-tRNA biosynthesis | Translation |
| 1806 | CCA tRNA nucleotidyltransferase (EC 2.7.7.72) | RNA transport | Translation |
| 157 | Transcription termination factor Rho | RNA degradation | Folding, sorting and degradation |
| 982 | RNA-binding protein Hfq | RNA degradation | Folding, sorting and degradation |
| 308 | Single-stranded DNA-binding protein | DNA replication | Replication and repair |
| 308 | Single-stranded DNA-binding protein | Mismatch repair | Replication and repair |
| 308 | Single-stranded DNA-binding protein | Homologous recombination | Replication and repair |
| 1930 | Exodeoxyribonuclease V gamma chain (EC 3.1.11.5) | Homologous recombination | Replication and repair |
| 1038 | Phospholipid ABC transporter substrate-binding protein MlaD | ABC transporters | Membrane transport |
| 1037 | Phospholipid ABC transporter permease protein MlaE | ABC transporters | Membrane transport |
| 1130 | Arginine ABC transporter%2C substrate-binding protein ArtI | ABC transporters | Membrane transport |
| 1131 | Arginine ABC transporter%2C permease protein ArtQ | ABC transporters | Membrane transport |
| 1742 | Peptide transport periplasmic protein sapA (TC 3.A.1.5.5) | ABC transporters | Membrane transport |
| 1743 | Peptide transport system permease protein sapB (TC 3.A.1.5.5) | ABC transporters | Membrane transport |
| 1744 | ABC transporter permease protein 2 (cluster 5%2C nickel/peptides/opines) | ABC transporters | Membrane transport |
| 1958 | Phosphoenolpyruvate-protein phosphotransferase of PTS system (EC 2.7.3.9) | Phosphotransferase system (PTS) | Membrane transport |
| 1957 | PTS system%2C glucose-specific IIA component (EC 2.7.1.199) | Phosphotransferase system (PTS) | Membrane transport |
| 1204 | PTS IIA-like nitrogen-regulatory protein PtsN | Phosphotransferase system (PTS) | Membrane transport |
| 1583 | Sensory histidine kinase QseC | Two-component system | Signal transduction |
| 858 | Fumarate reductase flavoprotein subunit (EC 1.3.5.4) | Two-component system | Signal transduction |
| 857 | Fumarate reductase iron-sulfur protein (EC 1.3.5.4) | Two-component system | Signal transduction |
| 855 | Fumarate reductase subunit D | Two-component system | Signal transduction |
| 103 | Nitrogen regulatory protein P-II | Two-component system | Signal transduction |
| 462 | ATP-dependent protease La (EC 3.4.21.53) Type I | Cell cycle - Caulobacter | Cell growth and death |
| 1155 | UDP-N-acetylglucosamine--N-acetylmuramyl-(pentapeptide) | Cell cycle - Caulobacter | Cell growth and death |
| 982 | RNA-binding protein Hfq | Quorum sensing | Cellular community - prokaryotes |
| 1365 | 2-keto-3-deoxy-D-arabino-heptulosonate-7-phosphate synthase I alpha (EC 2.5.1.54) | Quorum sensing | Cellular community - prokaryotes |
| 1583 | Sensory histidine kinase QseC | Quorum sensing | Cellular community - prokaryotes |
| 982 | RNA-binding protein Hfq | Biofilm formation - Vibrio cholerae | Cellular community - prokaryotes |
| 477 | Serine acetyltransferase (EC 2.3.1.30) | Biofilm formation - Vibrio cholerae | Cellular community - prokaryotes |
| 1957 | PTS system glucose-specific IIA component (EC 2.7.1.199) | Biofilm formation - Vibrio cholerae | Cellular community - prokaryotes |
| 1957 | PTS system glucose-specific IIA component (EC 2.7.1.199) | Biofilm formation - Escherichia coli | Cellular community - prokaryotes |
| 82 | Uncharacterized protease YegQ | Epithelial cell signaling in Helicobacter pylori infection | Other |
| 77 | AmpG permease | beta-Lactam resistance | Drug resistance: antimicrobial |
| 914 | CzcABC family efflux RND transporter%2C membrane fusion protein | beta-Lactam resistance | Drug resistance: antimicrobial |
| 915 | RND efflux system%2C inner membrane transporter | beta-Lactam resistance | Drug resistance: antimicrobial |
| 1149 | Cell division protein FtsI [Peptidoglycan synthetase] (EC 2.4.1.129) | beta-Lactam resistance | Drug resistance: antimicrobial |
| 1155 | UDP-N-acetylglucosamine--N-acetylmuramyl-(pentapeptide) | Vancomycin resistance | Drug resistance: antimicrobial |
| 1742 | Peptide transport periplasmic protein sapA (TC 3.A.1.5.5) | Cationic antimicrobial peptide (CAMP) resistance | Drug resistance: antimicrobial |
| 1743 | Peptide transport system permease protein sapB (TC 3.A.1.5.5) | Cationic antimicrobial peptide (CAMP) resistance | Drug resistance: antimicrobial |
| 1744 | ABC transporter%2C permease protein 2 (cluster 5%2C nickel/peptides/opines) | Cationic antimicrobial peptide (CAMP) resistance | Drug resistance: antimicrobial |
| 914 | CzcABC family efflux RND transporter membrane fusion protein | Cationic antimicrobial peptide (CAMP) resistance | Drug resistance: antimicrobial |
| 915 | RND efflux system%2C inner membrane transporter | Cationic antimicrobial peptide (CAMP) resistance | Drug resistance: antimicrobial |
| 932 | Acyl-[acyl-carrier-protein]--UDP-N-acetylglucosamine O-acyltransferase (EC 2.3.1.129) | Cationic antimicrobial peptide (CAMP) resistance | Drug resistance: antimicrobial |
| 1583 | Sensory histidine kinase QseC | Protein kinases | Protein families: metabolism |
| 1436 | Cell division-associated%2C ATP-dependent zinc metalloprotease FtsH | Peptidases and inhibitors | Protein families: metabolism |
| 462 | ATP-dependent protease La (EC 3.4.21.53) Type I | Peptidases and inhibitors | Protein families: metabolism |
| 82 | Uncharacterized protease YegQ | Peptidases and inhibitors | Protein families: metabolism |
| 339 | ADP-heptose--lipooligosaccharide heptosyltransferase II | Glycosyltransferases | Protein families: metabolism |
| 932 | Acyl-[acyl-carrier-protein]--UDP-N-acetylglucosamine O-acyltransferase (EC 2.3.1.129) | Lipopolysaccharide biosynthesis proteins | Protein families: metabolism |
| 339 | ADP-heptose--lipooligosaccharide heptosyltransferase II | Lipopolysaccharide biosynthesis proteins | Protein families: metabolism |
| 1783 | D-arabinose-5-phosphate isomerase (EC 5.3.1.13) | Lipopolysaccharide biosynthesis proteins | Protein families: metabolism |
| 1128 | D-sedoheptulose 7-phosphate isomerase (EC 5.3.1.28) | Lipopolysaccharide biosynthesis proteins | Protein families: metabolism |
| 1851 | D-glycero-beta-D-manno-heptose 1-phosphate adenylyltransferase (EC 2.7.7.70) | Lipopolysaccharide biosynthesis proteins | Protein families: metabolism |
| 1155 | UDP-N-acetylglucosamine--N-acetylmuramyl-(pentapeptide) | Peptidoglycan biosynthesis and degradation | Protein families: metabolism |
| 1149 | Cell division protein FtsI [Peptidoglycan synthetase] (EC 2.4.1.129) | Peptidoglycan biosynthesis and degradation | Protein families: metabolism |
| 51 | 1%2C6-anhydro-N-acetylmuramyl-L-alanine amidase | Peptidoglycan biosynthesis and degradation | Protein families: metabolism |
| 1965 | tRNA dimethylallyltransferase (EC 2.5.1.75) | Prenyltransferases | Protein families: metabolism |
| 1602 | Glutaminyl-tRNA synthetase (EC 6.1.1.18) | Amino acid related enzymes | Protein families: metabolism |
| 1420 | Glutamate-1-semialdehyde 2%2C1-aminomutase (EC 5.4.3.8) | Amino acid related enzymes | Protein families: metabolism |
| 534 | ATP synthase beta chain (EC 3.6.3.14) | Photosynthesis proteins | Protein families: metabolism |
| 533 | ATP synthase gamma chain (EC 3.6.3.14) | Photosynthesis proteins | Protein families: metabolism |
| 532 | ATP synthase alpha chain (EC 3.6.3.14) | Photosynthesis proteins | Protein families: metabolism |
| 531 | ATP synthase delta chain (EC 3.6.3.14) | Photosynthesis proteins | Protein families: metabolism |
| 529 | ATP synthase F0 sector subunit c (EC 3.6.3.14) | Photosynthesis proteins | Protein families: metabolism |
| 528 | ATP synthase F0 sector subunit a (EC 3.6.3.14) | Photosynthesis proteins | Protein families: metabolism |
| 527 | Permease of the drug/metabolite transporter (DMT) superfamily | Photosynthesis proteins | Protein families: metabolism |
| 1881 | Ferric uptake regulation protein FUR | Transcription factors | Genetic information processing |
| 1593 | Transcription termination protein NusB | Transcription machinery | Genetic information processing |
| 157 | Transcription termination factor Rho | Transcription machinery | Genetic information processing |
| 157 | Transcription termination factor Rho | Messenger RNA biogenesis | Genetic information processing |
| 982 | RNA-binding protein Hfq | Messenger RNA biogenesis | Genetic information processing |
| 385 | LSU ribosomal protein L17p | Ribosome | Genetic information processing |
| 1593 | Transcription termination protein NusB | Ribosome biogenesis | Genetic information processing |
| 379 | ATP-dependent RNA helicase SrmB | Ribosome biogenesis | Genetic information processing |
| 1806 | CCA tRNA nucleotidyltransferase (EC 2.7.7.72) | Transfer RNA biogenesis | Genetic information processing |
| 1602 | Glutaminyl-tRNA synthetase (EC 6.1.1.18) | Transfer RNA biogenesis | Genetic information processing |
| 1965 | tRNA dimethylallyltransferase (EC 2.5.1.75) | Transfer RNA biogenesis | Genetic information processing |
| 2029 | tRNA-5-carboxymethylaminomethyl-2-thiouridine(34) synthesis protein MnmE | Transfer RNA biogenesis | Genetic information processing |
| 523 | tRNA-5-carboxymethylaminomethyl-2-thiouridine(34) synthesis protein MnmG | Transfer RNA biogenesis | Genetic information processing |
| 644 | Translation elongation factor G | Translation factors | Genetic information processing |
| 1920 | Chaperone protein DnaJ | Chaperones and folding catalysts | Genetic information processing |
| 1436 | Cell division-associated%2C ATP-dependent zinc metalloprotease FtsH | Chaperones and folding catalysts | Genetic information processing |
| 528 | ATP synthase F0 sector subunit a (EC 3.6.3.14) | Chaperones and folding catalysts | Genetic information processing |
| 308 | Single-stranded DNA-binding protein | DNA replication proteins | Genetic information processing |
| 1354 | DNA-binding protein H-NS | Chromosome and associated proteins | Genetic information processing |
| 982 | RNA-binding protein Hfq | Chromosome and associated proteins | Genetic information processing |
| 1149 | Cell division protein FtsI [Peptidoglycan synthetase] (EC 2.4.1.129) | Chromosome and associated proteins | Genetic information processing |
| 914 | CzcABC family efflux RND transporter%2C membrane fusion protein | Chromosome and associated proteins | Genetic information processing |
| 60 | Rod shape-determining protein MreD | Chromosome and associated proteins | Genetic information processing |
| 523 | tRNA-5-carboxymethylaminomethyl-2-thiouridine(34) synthesis protein MnmG | Chromosome and associated proteins | Genetic information processing |
| 629 | Site-specific tyrosine recombinase XerC | Chromosome and associated proteins | Genetic information processing |
| 249 | Site-specific tyrosine recombinase XerD | Chromosome and associated proteins | Genetic information processing |
| 308 | Single-stranded DNA-binding protein | DNA repair and recombination proteins | Genetic information processing |
| 1930 | Exodeoxyribonuclease V gamma chain (EC 3.1.11.5) | DNA repair and recombination proteins | Genetic information processing |
| 1354 | DNA-binding protein H-NS | DNA repair and recombination proteins | Genetic information processing |
| 644 | Translation elongation factor G | Mitochondrial biogenesis | Genetic information processing |
| 308 | Single-stranded DNA-binding protein | Mitochondrial biogenesis | Genetic information processing |
| 1920 | Chaperone protein DnaJ | Mitochondrial biogenesis | Genetic information processing |
| 1038 | Phospholipid ABC transporter substrate-binding protein MlaD | Transporters | Signaling and cellular processes |
| 1037 | Phospholipid ABC transporter permease protein MlaE | Transporters | Signaling and cellular processes |
| 1130 | Arginine ABC transporter%2C substrate-binding protein ArtI | Transporters | Signaling and cellular processes |
| 1131 | Arginine ABC transporter%2C permease protein ArtQ | Transporters | Signaling and cellular processes |
| 1742 | Peptide transport periplasmic protein sapA (TC 3.A.1.5.5) | Transporters | Signaling and cellular processes |
| 1743 | Peptide transport system permease protein sapB (TC 3.A.1.5.5) | Transporters | Signaling and cellular processes |
| 1744 | ABC transporter%2C permease protein 2 (cluster 5%2C nickel/peptides/opines) | Transporters | Signaling and cellular processes |
| 77 | AmpG permease | Transporters | Signaling and cellular processes |
| 1958 | Phosphoenolpyruvate-protein phosphotransferase of PTS system (EC 2.7.3.9) | Transporters | Signaling and cellular processes |
| 1957 | PTS system%2C glucose-specific IIA component (EC 2.7.1.199) | Transporters | Signaling and cellular processes |
| 1204 | PTS IIA-like nitrogen-regulatory protein PtsN | Transporters | Signaling and cellular processes |
| 1009 | Tol-Pal system protein TolQ | Transporters | Signaling and cellular processes |
| 1010 | Tol biopolymer transport system%2C TolR protein | Transporters | Signaling and cellular processes |
| 488 | Inner membrane component of TAM transport system | Transporters | Signaling and cellular processes |
| 915 | RND efflux system%2C inner membrane transporter | Transporters | Signaling and cellular processes |
| 812 | Pantothenate:Na+ symporter (TC 2.A.21.1.1) | Transporters | Signaling and cellular processes |
| 1302 | Sodium/glutamate symporter | Transporters | Signaling and cellular processes |
| 1355 | Na+/H+ antiporter | Transporters | Signaling and cellular processes |
| 1013 | Tol-Pal system peptidoglycan-associated lipoprotein PAL | Transporters | Signaling and cellular processes |
| 1012 | Tol-Pal system beta propeller repeat protein TolB | Transporters | Signaling and cellular processes |
| 914 | CzcABC family efflux RND transporter%2C membrane fusion protein | Transporters | Signaling and cellular processes |
| 1583 | Sensory histidine kinase QseC | Two-component system | Signaling and cellular processes |
| 1691 | Glucose-6-phosphate isomerase (EC 5.3.1.9) | Exosome | Signaling and cellular processes |
| 572 | Succinate dehydrogenase flavin-adding protein%2C antitoxin of CptAB toxin-antitoxin | Prokaryotic defense system | Signaling and cellular processes |
| 1355 | Na+/H+ antiporter | Antimicrobial resistance genes | Signaling and cellular processes |
| 914 | CzcABC family efflux RND transporter%2C membrane fusion protein | Antimicrobial resistance genes | Signaling and cellular processes |
| 915 | RND efflux system%2C inner membrane transporter | Antimicrobial resistance genes | Signaling and cellular processes |
| 1505 | Bacterial non-heme ferritin (EC 1.16.3.2) | Enzymes with EC numbers | Unclassified: metabolism |
| 1035 | Apolipoprotein N-acyltransferase / Copper homeostasis protein CutE | Enzymes with EC numbers | Unclassified: metabolism |
| 180 | FAD:protein FMN transferase (EC 2.7.1.180) | Enzymes with EC numbers | Unclassified: metabolism |
| 186 | Na(+)-translocating NADH-quinone reductase subunit A (EC 1.6.5.8) | Enzymes with EC numbers | Unclassified: metabolism |
| 185 | Na(+)-translocating NADH-quinone reductase subunit B (EC 1.6.5.8) | Enzymes with EC numbers | Unclassified: metabolism |
| 184 | Na(+)-translocating NADH-quinone reductase subunit C (EC 1.6.5.8) | Enzymes with EC numbers | Unclassified: metabolism |
| 183 | Na(+)-translocating NADH-quinone reductase subunit D (EC 1.6.5.8) | Enzymes with EC numbers | Unclassified: metabolism |
| 181 | Na(+)-translocating NADH-quinone reductase subunit F (EC 1.6.5.8) | Enzymes with EC numbers | Unclassified: metabolism |
| 1712 | probable membrane protein YPO2224 | Glycan metabolism | Unclassified: metabolism |
| 1394 | LPS-core synthesis glycosyltransferase PM0509 | Glycan metabolism | Unclassified: metabolism |
| 607 | Uncharacterized protein EC-HemY in Proteobacteria | Cofactor metabolism | Unclassified: metabolism |
| 349 | Murein hydrolase activator NlpD | Structural proteins | Unclassified: signaling and cellular processes |
| 257 | Cell envelope opacity-associated protein A | Structural proteins | Unclassified: signaling and cellular processes |
| 2014 | Cytoskeleton protein RodZ | Structural proteins | Unclassified: signaling and cellular processes |
| 552 | UPF0438 protein YifE | Function unknown | Poorly characterized |
| 1803 | Phosphate transport regulator (distant homolog of PhoU) | Function unknown | Poorly characterized |

**Table S5A.** RNAseq analysis of *Hp* 392 comparing anaerobic to aerobic conditions showing genes that were significantly upregulated (≥ 2-fold) anaerobically.

| **PEG** | **Annotation** | **Fold Change** |
| --- | --- | --- |
| 1506 | hypothetical protein | 23.46 |
| 593 | 16S rRNA (guanine(1516)-N(2))-methyltransferase | 9.66 |
| 620 | Uncharacterized MFS-type transporter | 9.62 |
| 592 | Sigma factor RpoE regulatory protein RseC | 8.64 |
| 826 | Transcriptional regulator%2C MerR family | 6.52 |
| 619 | Molybdopterin-guanine dinucleotide biosynthesis protein MobB | 6.05 |
| 1961 | hypothetical protein | 5.98 |
| 989 | Uncharacterized transporter YfbS | 5.67 |
| 824 | Dihydroneopterin aldolase | 5.58 |
| 1012 | Monofunctional biosynthetic peptidoglycan transglycosylase | 5.41 |
| 1505 | hypothetical protein | 4.54 |
| 191 | UPF0235 protein VC0458 | 4.52 |
| 1229 | Protein SprT | 4.28 |
| 639 | hypothetical protein | 4.28 |
| 594 | tRNA (uracil(54)-C5)-methyltransferase | 4.27 |
| 1032 | Iron compound ABC uptake transporter permease protein | 4.22 |
| 637 | tRNA 5-methylaminomethyl-2-thiouridine synthase subunit TusD | 3.78 |
| 1621 | tRNA pseudouridine(55) synthase | 3.77 |
| 19 | Formamidopyrimidine-DNA glycosylase | 3.72 |
| 1013 | Transcriptional repressor protein TrpR | 3.68 |
| 78 | AmpG permease | 3.39 |
| 1768 | Hypothetical metal-binding enzyme%2C YcbL homolog | 3.33 |
| 510 | Lipid A export permease/ATP-binding protein MsbA | 3.32 |
| 664 | Transcriptional regulator IlvY%2C LysR family | 3.25 |
| 638 | tRNA 5-methylaminomethyl-2-thiouridine synthase subunit TusC | 3.24 |
| 710 | hypothetical protein | 3.18 |
| 1014 | Soluble lytic murein transglycosylase (EC 4.2.2.n1) | 3.17 |
| 509 | ABC-type multidrug transport system%2C ATPase | 3.15 |
| 988 | Uncharacterized transporter YfbS | 3.1 |
| 711 | Phosphoribosylformylglycinamidine synthase%2C synthetase subunit | 3.09 |
| 1620 | Chorismate mutase I (EC 5.4.99.5) / Cyclohexadienyl dehydrogenase | 3.07 |
| 368 | Iron(III) dicitrate transport ATP-binding protein FecE | 3.02 |
| 645 | hypothetical protein | 3.02 |
| 837 | Transcriptional regulator of glmS gene%2C DeoR family | 2.96 |
| 536 | hypothetical protein | 2.95 |
| 389 | tRNA pseudouridine(13) synthase | 2.95 |
| 1276 | 2-amino-4-hydroxy-6-hydroxymethyldihydropteridine pyrophosphokinase | 2.86 |
| 987 | Ribonuclease HII (EC 3.1.26.4) | 2.8 |
| 1625 | Bacterial ribosome SSU maturation protein RimP | 2.8 |
| 304 | hypothetical protein | 2.8 |
| 1685 | Phage integrase | 2.76 |
| 618 | Sigma factor RpoE negative regulatory protein RseB precursor | 2.76 |
| 986 | Lipid-A-disaccharide synthase (EC 2.4.1.182) | 2.74 |
| 636 | YheO-like PAS domain | 2.71 |
| 1928 | Uncharacterized protein YdiJ | 2.7 |
| 508 | Ferric vulnibactin receptor VuuA | 2.69 |
| 1277 | Poly(A) polymerase (EC 2.7.7.19) | 2.65 |
| 386 | Cobalamin biosynthesis protein CobN and related Mg-chelatases | 2.65 |
| 1031 | Iron compound ABC uptake transporter permease protein | 2.64 |
| 1622 | Ribosome-binding factor A | 2.54 |
| 1102 | UDP-N-acetylglucosamine 1-carboxyvinyltransferase | 2.53 |
| 1624 | Transcription termination protein NusA | 2.52 |
| 1935 | hypothetical protein | 2.51 |
| 160 | Transcription termination factor Rho | 2.5 |
| 367 | Iron(III) dicitrate transport system permease protein FecD | 2.47 |
| 146 | Smp-like protein | 2.46 |
| 1326 | hypothetical protein | 2.46 |
| 1429 | LPS-core synthesis glycosyltransferase PM0509 | 2.43 |
| 843 | UPF0701 protein YicC | 2.41 |
| 1237 | RNase adapter protein RapZ | 2.37 |
| 1623 | Translation initiation factor 2 | 2.36 |
| 1697 | hypothetical protein | 2.36 |
| 1000 | Methionine repressor MetJ | 2.33 |
| 1486 | Probable endopeptidase NlpC | 2.32 |
| 597 | Protein YihD | 2.29 |
| 1426 | dsDNA mimic protein%2C highly acidic | 2.29 |
| 157 | Transposase | 2.28 |
| 1696 | putative membrane protein | 2.28 |
| 74 | Octanoate-[acyl-carrier-protein]-protein-N-octanoyltransferase | 2.25 |
| 818 | Uncharacterized MFS-type transporter | 2.24 |
| 349 | hypothetical protein | 2.23 |
| 388 | 5'-nucleotidase SurE | 2.19 |
| 71 | Septum-associated rare lipoprotein A | 2.18 |
| 140 | Membrane-bound lytic murein transglycosylase F | 2.18 |
| 513 | converved hypothetical protein | 2.17 |
| 1670 | 2-oxoglutarate dehydrogenase E1 component | 2.14 |
| 1947 | DNA mismatch repair protein MutL | 2.13 |
| 227 | FIG021862: membrane protein%2C exporter | 2.12 |
| 1421 | Transcriptional regulator%2C MerR family | 2.12 |
| 836 | Glutamine--fructose-6-phosphate aminotransferase [isomerizing] | 2.12 |
| 1249 | Acyl-CoA thioesterase II | 2.12 |
| 948 | hypothetical protein | 2.11 |
| 1358 | Lipopolysaccharide export system permease protein LptG | 2.11 |
| 889 | Putative sulfate permease | 2.11 |
| 1774 | Replication-associated recombination protein RarA | 2.1 |
| 1767 | FIG001587: exported protein | 2.1 |
| 598 | Molybdenum cofactor guanylyltransferase | 2.1 |
| 1198 | Protein PhnA | 2.09 |
| 1096 | Phospholipid ABC transporter ATP-binding protein MlaF | 2.08 |
| 76 | hypothetical protein | 2.08 |
| 1752 | Peptide chain release factor N(5)-glutamine methyltransferase | 2.07 |
| 281 | Diacylglycerol kinase | 2.05 |
| 607 | DNA polymerase III delta prime subunit | 2.03 |
| 1700 | Translation initiation factor SUI1-related protein | 2.03 |
| 1030 | Murein DD-endopeptidase MepM | 2.03 |
| 565 | Flavoprotein MioC | 2.02 |
| 1189 | UDP-N-acetylglucosamine--N-acetylmuramyl-(pentapeptide) | 2.02 |
| 1191 | D-alanine--D-alanine ligase | 2.01 |

**Table S5B.** RNAseq analysis of *Hp* 392 comparing anaerobic to aerobic conditions showing genes that were significantly downpregulated (≥ 2-fold) anaerobically.

| **PEG** | **Annotation** | **Fold Change** |
| --- | --- | --- |
| 1566 | TonB-dependent receptor%3B Outer membrane receptor for ferrienterochelin and colicins | -8.2 |
| 1174 | NAD-dependent protein deacetylase of SIR2 family | -4.58 |
| 1820 | ABC transporter%2C ATP-binding protein | -4.51 |
| 845 | L-lactate permease | -4.4 |
| 142 | Ribosome-associated inhibitor A | -4.3 |
| 949 | FKBP-type peptidyl-prolyl cis-trans isomerase FklB | -4.23 |
| 1827 | TonB-dependent receptor | -4.18 |
| 700 | Sodium-Choline Symporter | -3.94 |
| 1826 | hypothetical protein | -3.84 |
| 352 | putative phage shock protein E precursor | -3.5 |
| 458 | hypothetical protein | -3.49 |
| 1087 | Galactose/methyl galactoside ABC transporter%2C substrate-binding protein MglB | -3.32 |
| 1825 | Vitamin B12 ABC transporter%2C substrate-binding protein | -3.29 |
| 1088 | Galactose operon repressor%2C GalR-LacI family of transcriptional regulators | -3.24 |
| 1561 | ABC transporter%2C substrate-binding protein (cluster 8%2C B12/iron complex) | -3.19 |
| 1046 | hypothetical protein | -3.19 |
| 95 | Chaperone protein ClpB (ATP-dependent unfoldase) | -3.15 |
| 539 | thiamine-phosphate pyrophosphorylase | -3.08 |
| 1092 | Aldose 1-epimerase | -2.98 |
| 1091 | Galactokinase | -2.98 |
| 1085 | Galactose/methyl galactoside ABC transporter%2C permease protein MglC | -2.92 |
| 1824 | ABC transporter%2C permease protein (cluster 8%2C B12/iron complex) | -2.9 |
| 45 | Autonomous glycyl radical cofactor | -2.77 |
| 1064 | Cytochrome d ubiquinol oxidase subunit I | -2.77 |
| 243 | Carboxymuconolactone decarboxylase family protein | -2.74 |
| 1065 | Cytochrome d ubiquinol oxidase subunit II | -2.72 |
| 1170 | putative 5'(3')-deoxyribonucleotidase | -2.65 |
| 1900 | Chaperone protein DnaK | -2.61 |
| 891 | NADH dehydrogenase (EC 1.6.99.3) | -2.58 |
| 1672 | Putative TEGT family carrier/transport protein | -2.58 |
| 555 | 2-oxoglutarate/malate translocator | -2.57 |
| 1090 | Galactokinase (EC 2.7.1.6) | -2.53 |
| 1086 | Galactose/methyl galactoside ABC transporter%2C ATP-binding protein MglA | -2.42 |
| 1564 | TonB-dependent receptor%3B Outer membrane receptor for ferrienterochelin and colicins | -2.37 |
| 801 | Methionine ABC transporter ATP-binding protein | -2.32 |
| 308 | hypothetical protein | -2.32 |
| 1575 | DNA protection during starvation protein | -2.31 |
| 1048 | Histidine permease YuiF | -2.29 |
| 108 | Serine hydroxymethyltransferase | -2.28 |
| 991 | Aspartate ammonia-lyase | -2.27 |
| 504 | ATP-dependent protease | -2.25 |
| 1954 | Guanylate kinase | -2.21 |
| 1307 | Ferric iron ABC transporter%2C iron-binding protein | -2.21 |
| 1206 | FIG024746: hypothetical protein | -2.19 |
| 52 | 1%2C6-anhydro-N-acetylmuramyl-L-alanine amidase | -2.19 |
| 442 | LSU ribosomal protein L29p (L35e) | -2.16 |
| 724 | Spermidine/putrescine import ABC transporter substrate-binding protein PotD | -2.15 |
| 950 | Phosphatidylserine decarboxylase | -2.15 |
| 467 | Hybrid peroxiredoxin hyPrx5 | -2.14 |
| 441 | SSU ribosomal protein S17p (S11e) | -2.13 |
| 1089 | Galactose-1-phosphate uridylyltransferase | -2.13 |
| 1544 | Bacterial non-heme ferritin | -2.12 |
| 1533 | ABC-type Co2+ transport system%2C periplasmic component | -2.1 |
| 880 | Di- and tricarboxylate transporters | -2.1 |
| 1907 | Molybdenum ABC transporter%2C substrate-binding protein ModA | -2.1 |
| 1732 | Uncharacterized integral membrane protein GSU2901 | -2.1 |
| 289 | hypothetical protein | -2.07 |
| 998 | Carbon storage regulator | -2.06 |
| 1822 | Molybdenum ABC transporter%2C substrate-binding protein ModA | -2.06 |
| 1882 | FIG005121: SAM-dependent methyltransferase | -2.06 |
| 373 | Phosphoenolpyruvate carboxykinase [ATP] | -2.06 |
| 1796 | CRISPR-associated endonuclease Cas9 | -2.05 |
| 761 | RNA polymerase sigma factor RpoD | -2.05 |
| 993 | Heat shock protein 60 kDa family chaperone GroEL | -2.03 |
| 1705 | SSU ribosomal protein S1p | -2.02 |
| 1894 | Dihydrolipoamide dehydrogenase of pyruvate dehydrogenase complex | -2.02 |
| 1162 | Thioredoxin | -2.02 |
| 1716 | mercuric ion transport protein | -2.01 |
| 1380 | UPF0033 protein YedF | -2 |

**Table S6**. Strains of *Haemophilus parainfluenzae* used to align the essential genome of *Hp* 392 and EL1 to identify the core essential genome of *Hp*.

| **NCBI RefSeq** | **Genome assembly** | **Strain** | **Taxon** |
| --- | --- | --- | --- |
| GCF_000210895.1 | ASM21089v1 | T3T1 | *Haemophilus parainfluenzae* |
| GCF 003390455 1 | ASM339045v1 | M27794 | *Haemophilus parainfluenzae* |
| GCF 004104465 2 | ASM410446v2 | LC_1315_18 | *Haemophilus parainfluenzae* |
| GCF 014931275 1 | ASM1493127v1 | M1C160_1 | *Haemophilus parainfluenzae* |
| GCF 014931295 1 | ASM1493129v1 | M1C152_1 | *Haemophilus parainfluenzae* |
| GCF 014931315 1 | ASM1493131v1 | M1C149_1 | *Haemophilus parainfluenzae* |
| GCF 014931335 1 | ASM1493133v1 | M1C147_1 | *Haemophilus parainfluenzae* |
| GCF 014931355 1 | ASM1493135v1 | M1C146_1 | *Haemophilus parainfluenzae* |
| GCF 014931375 1 | ASM1493137v1 | M1C142_1 | *Haemophilus parainfluenzae* |
| GCF 014931395 1 | ASM1493139v1 | M1C137_2 | *Haemophilus parainfluenzae* |
| GCF 014931415 1 | ASM1493141v1 | M1C130_2 | *Haemophilus parainfluenzae* |
| GCF 014931435 1 | ASM1493143v1 | M1C125_4 | *Haemophilus parainfluenzae* |
| GCF 014931455 1 | ASM1493145v1 | M1C120_2 | *Haemophilus parainfluenzae* |
| GCF 014931475 1 | ASM1493147v1 | M1C113_1 | *Haemophilus parainfluenzae* |
| GCF 016127215 1 | ASM1612721v1 | FDAARGOS_1000 | *Haemophilus parainfluenzae* |
| GCF 900638025 1 | 55312 D01 | NCTC10665 | *Haemophilus parainfluenzae* |

See separate file for Table S7

**Table S7.** Essential genes for *Hp* 392 and *Hp* EL1. PEG “392” indicates the PEG number in *Hp* 392 and PEG “EL1” the PEG number in the respective homologue gene in *Hp* EL1. The highest identity score (not shown) was considered to identify the homologues when one gene aligned to two or more genes. “Absolute” indicates genes essentiality *in vitro* (genes that are part of the essential genome); “Aerobic” and “Anaerobic” indicates conditionally essential gene for survival in biofilm aerobically and anaerobically, respectively; “Reads” indicates average number of transposon reads in each gene; and “Essentiality” indicates if the gene is essential (Y) or not essential (N) in each condition tested.
